# Species abundances often conform to ‘abundant-centre’ patterns depending on dispersal capabilities

**DOI:** 10.1101/2023.03.31.535106

**Authors:** Connor T. Panter, Steven P. Bachman, Oliver Baines, Helge Bruelheide, Stephan Kambach, Maria Sporbert, Richard Field, Franziska Schrodt

## Abstract

A shared goal within macroecology, biogeography and population ecology research is to understand biodiversity patterns and the processes driving them across spatial and taxonomic scales. A common approach to study macroecological patterns and processes involves developing and testing ecogeographical rules or hypotheses. The much-debated ‘abundant-centre’ hypothesis posits that species’ abundances are highest in their range centres and decline towards their range edges. We perform the largest global test of the hypothesis to date, on 3,675 species, using 6,055,549 abundance observations. Using meta-analytical approaches, we summarised species-level abundance–distance correlations exploring the effects of dispersal-related species traits on abundance–distance relationships. Overall, animals did not follow abundant-centre patterns, whereas plants tended to. Larger-bodied mammals were more likely to conform to abundant-centre patterns, as were mammals and freshwater fishes from higher latitudes. Perennial life cycles and large range sizes were significant predictors of abundant-centre patterns in plants. Trees and shrubs with larger seeds showed more support for abundant-centre patterns. Accounting for species dispersal improves models of abundant-centre patterns across geographic space. Assuming abundant-centre patterns represent optimal equilibria within nature, our findings suggest that abundant-centre relationships are not a general ecological phenomenon but tend to manifest only in species with higher dispersal capabilities.

## MAIN TEXT

Understanding patterns of biodiversity and the processes that drive them across varying spatial, temporal and taxonomic scales is a shared goal within macroecology, biogeography and population ecology research (Beck *et al*. 2012; Shade *et al*. 2018; McGill 2018). Such information is essential to improving our understanding of species’ responses to environmental change, and can be applied to invasive species management (Hui *et al*. 2011; Hansen *et al*. 2013) and conservation planning (Pearce & Ferrier 2001; Santini *et al*. 2021).

A common approach to studying macroecological patterns and processes is to develop and test ecogeographical “rules” or “hypotheses”. As outlined by Baiser *et al*. (2019), over the last decade, the study of ecogeographical rules has seen a resurgence due to increased data availability, methodological advancements and the need for applied research efforts, e.g., in invasive species management (Lomolino *et al*. 2006; Mathys & Lockwood 2009; Hansen *et al*. 2013) and environmental change research (Millien *et al*. 2006; Gardner *et al*. 2011; Ehrlén & Morris 2015). Historically, many ecogeographical rules were developed to describe scaling relationships in species’ diversity (Pianka 1966), body mass (Bergmann 1847), range size (Stevens 1989, 1992) and geographical distributions of populations (Grinnell 1922), such as the ‘abundant-centre’ hypothesis.

The abundant-centre hypothesis derives from early ideas proposed by Grinnell (1922), who likened the distribution and dispersal of animal populations to those of gas molecules in space (Santini *et al*. 2018; see Pironon *et al*. 2017 for an overview). The abundant-centre hypothesis posits that species abundances are highest in their range centres and decline towards their range edges (Sagarin & Gaines 2002a; McGill & Collins 2003; Sexton *et al*. 2009; Rivadeneira *et al*. 2010; Pironon *et al*. 2016; Santini *et al*. 2018; Shalom *et al*. 2020; Dallas *et al*. 2017, 2020). The hypothesis assumes that 1) a species’ geographic range is a representation of its environmental or ecological niche (Hutchinson 1957) and 2) that environmental conditions are more optimal near the centre of a species’ range and harsher towards the range edges (Brown 1984; Pironon *et al*. 2016). Since its inception, the abundant-centre hypothesis has been debated by macroecologists and biogeographers alike (see Osorio-Olvera *et al*. 2016; Dallas *et al*. 2017; Soberón *et al*. 2018), and has been tested within multiple taxonomic groups (birds: Freeman & Beehler 2012; Burner *et al*. 2019; Osorio-Olvera *et al*. 2020; vascular plants: Pérez-Collazos *et al*. 2009; Dixon *et al*. 2013; Sporbert *et al*. 2020; reef fishes: Tuya *et al*. 2008; Waldock *et al*. 2019; Shalom *et al*. 2020; mammals: Virgós *et al*. 2011; Martínez-Gutiérrez *et al*. 2017; Wen *et al*. 2020; Chaiyes *et al*. 2020; coastal invertebrates: Sagarin & Gaines 2002b; Rivadeneira *et al*. 2010; Tam & Scrosati 2011; Baldanzi *et al*. 2013; Scrosati & Freeman 2019; Ntuli *et al*. 2020; as well as across multiple taxonomic groups: Dallas *et al*. 2017; Santini *et al*. 2018; Chevalier *et al*. 2021). These studies employed an array of methodological approaches, e.g., focusing on centrality (Martínez-Meyer *et al*. 2013, Dallas *et al*. 2017) as opposed to marginality (Blackburn *et al*. 1999; Sexton *et al*. 2009; see Santini *et al*. 2018).

Traditionally, the abundant-centre hypothesis has been tested across species’ geographic space (Sagarin & Gaines 2002a; Rivadeneira *et al*. 2010; Baldanzi *et al*. 2013; Shalom *et al*. 2020). Support for the hypothesis remains scarce and recent research has shifted focus towards exploring abundance distributions across species’ environmental or ecological niche space (VanDerWal *et al*. 2009; Martínez-Gutiérrez *et al*. 2017; Osorio-Olvera *et al*. 2019, 2020; Yáñez-Arenas *et al*. 2020; Chaiyes *et al*. 2020). For example, Dallas *et al*. (2017) conducted a test of the abundant-centre hypothesis on 1,400 species of North American birds, mammals, freshwater fishes and trees. The authors tested the hypothesis across both species’ geographic and ecological niche space and found no consistent relationships between abundance and distance from range centre (hereafter ‘abundance–distance relationships’) in either case (Dallas *et al*. 2017). In addition, Sporbert *et al*. (2020) tested multiple macroecological rules, including the abundant-centre hypothesis, on 517 species of European vascular plants, concluding mixed support probably driven by various environmental factors. A meta-analysis across a larger pool of taxa concluded that relationships between species’ abundances and environmental suitability are weak or absent, and can even be reversed (Weber *et al*. 2017). To date, such mixed support for the abundant-centre hypothesis appears to be repeatedly found across both geographic and ecological niche space (Dallas *et al*. 2017; Santini *et al*. 2018; Sporbert *et al*. 2020; Chevalier *et al*. 2021). It also remains unclear why abundant-centre patterns are rarely dominant (although see Hidas *et al*. 2010; Scrosati & Freeman 2019), but reasons may include differences in methodologies used (Santini *et al*. 2018), interpretation of results and/or taxonomic groups studied, e.g., species with unique life histories such as juvenile planktonic life stages (see Ntuli *et al*. 2020).

Assuming that an abundant-centre distribution represents a natural equilibrium of species populations within nature (as implied by Grinnell (1922)), optimal environments occur in the centre of a species’ range, and that the environment is spatially autocorrelated, we would expect to find that (i) species with higher dispersal capabilities, i.e., birds, larger-bodied mammals (e.g., large-bodied ungulates vs. small mammals) and plants with faster life cycles (e.g., annuals vs. perennials) tend to follow abundant-centre patterns as these species are better at tracking changing environments (Kokko & López-Sepulcre 2006; Thompson & Gonzalez 2017). These species are better at maintaining higher abundances in their range centres, where environmental conditions are assumed to be most favourable (Fristoe *et al*. 2022). In line with previous research, we expect to find that (ii) species with invasive tendencies, e.g., the Cane Toad (*Rhinella marina*), may not display abundant-centre distributions and may in fact conform to the opposite, i.e., abundances highest at range edges which act as invasion fronts (Trumbo *et al*. 2016). Here, we explored the effect of range size on abundance– distance relationships to account for the length of the range size gradient, i.e., abundant-centre patterns may be more detectable along longer-range size gradients than shorter gradients. Finally, species with larger ranges tend to have higher abundances than those with smaller ranges (Gaston *et al*. 2000), and may be able to cope with suboptimal environmental conditions better and thus maintain abundant-centre patterns.

Despite the recent shift towards testing the abundant-centre hypothesis across species’ ecological niche spaces, key research gaps remain regarding relationships in geographical space. Linking species’ ecological characteristics (traits), such as dispersal capabilities (Feng & Qiao 2022), to the species’ sampled abundances allows for a more sophisticated examination of the abundant-centre hypothesis using real-world data. Yet, other than Dallas *et al*. (2017) who explored the effect of body size on support for the abundant-centre hypothesis and Rivadeneira *et al*. (2010) who linked life history traits to porcelain crab abundance distributions, to date there have been no thorough examinations of the effects of species’ traits on support for the hypothesis across large taxonomic and spatial scales. By quantifying species’ traits, we may be able to explain a sizeable proportion of variation reported in previous findings.

Here, we use a trait-based approach to conduct the largest test of the abundant-centre hypothesis to date across five major taxonomic groups: birds, mammals, freshwater fishes, reef fishes and plants. Given that previous research has suggested that support for the hypothesis may be scale-dependent (Dallas *et al*. 2017; Santini *et al*. 2018), we also explore the effects of scale. We employ a meta-analytical approach to explore the effects of species’ traits and synthesise findings from previously published studies. Research has suggested that abundance patterns are closely related to species’ dispersal capabilities in particular (see Santini *et al*. 2018; Dallas & Santini 2020; Feng & Qiao 2022). We use real-world abundance data to explore the effects of spatial and taxonomic scale and species traits related to dispersal capabilities, for animals: 1) taxonomic group (categorical), 2) body size (cm; g), 3) invasiveness (binary), 4) feeding guild (categorical), 5) range size (km^2^), 6) absolute latitude (°), 7) extent (km), 8) grain (km^2^) and 9) focus (km^2^); see Methods and Materials for full descriptions of scale variables); and for plants: 1) functional group (categorical), 2) mean plant height (m), 3) invasiveness (binary), 4) life span (categorical), 5) life form (categorical), 6) seed mass (mg), 7) range size (km^2^), 8) absolute latitude (°), 9) extent (km), 10) grain (km^2^) and 11) focus (km^2^).

We ran a meta-analysis using abundance–distance correlation coefficients for 3,675 species (3,060 animal species; 615 plant species), representing 6,055,549 observations (3,698,959 animals; 2,356,590 plants) from 14 studies (Table 1; Fig. 1). Our data included 1,683 bird species, 1,131 reef fish species, 202 mammals species and 44 freshwater fish species. For plants the data set comprised 401 herb species, 120 tree species, 65 grass species and 29 shrub species. The mean number of abundance observations per animal species was 1,209 ± 1,944 (range: 5 – 11,948) and for plants was 3,832 ± 5,988 (range: 5 – 67,486) (Table 1). Our test of the abundant-centre hypothesis remains the largest to date, in terms of sample size and global coverage.

**Table 1.**
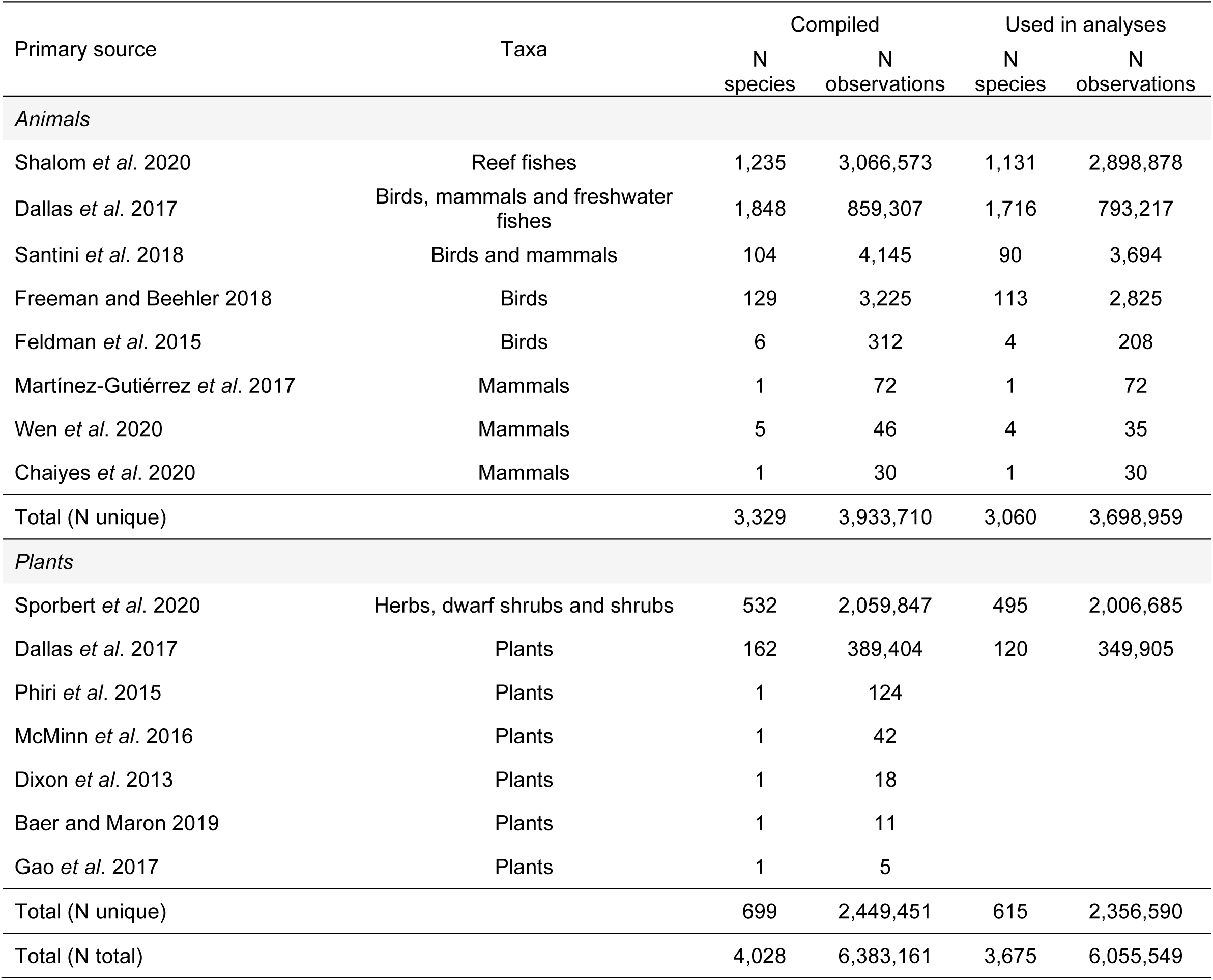
Overview of primary sources, taxonomic group, number of species and observations used to calculate abundance–distance correlations. Due to limited coverage of species trait data, some species were omitted prior to statistical analyses.

| Primary source | Taxa | Compiled |  | Used in analyses |  |
| --- | --- | --- | --- | --- | --- |
|  |  | N<br>species | N<br>observations | N<br>species | N<br>observations |
| Animals |  |  |  |  |  |
| Shalom *et al.* 2020 | Reef fishes | 1,235 | 3,066,573 | 1,131 | 2,898,878 |
| Dallas *et al.* 2017 | Birds, mammals and freshwater fishes | 1,848 | 859,307 | 1,716 | 793,217 |
| Santini *et al.* 2018 | Birds and mammals | 104 | 4,145 | 90 | 3,694 |
| Freeman and Beehler 2018 | Birds | 129 | 3,225 | 113 | 2,825 |
| Feldman *et al.* 2015 | Birds | 6 | 312 | 4 | 208 |
| Martínez-Gutiérrez *et al.* 2017 | Mammals | 1 | 72 | 1 | 72 |
| Wen *et al.* 2020 | Mammals | 5 | 46 | 4 | 35 |
| Chaiyes *et al.* 2020 | Mammals | 1 | 30 | 1 | 30 |
| Total (N unique) |  | 3,329 | 3,933,710 | 3,060 | 3,698,959 |
| Plants |  |  |  |  |  |
| Sporbert *et al.* 2020 | Herbs, dwarf shrubs and shrubs | 532 | 2,059,847 | 495 | 2,006,685 |
| Dallas *et al.* 2017 | Plants | 162 | 389,404 | 120 | 349,905 |
| Phiri *et al.* 2015 | Plants | 1 | 124 |  |  |
| McMinn *et al.* 2016 | Plants | 1 | 42 |  |  |
| Dixon *et al.* 2013 | Plants | 1 | 18 |  |  |
| Baer and Maron 2019 | Plants | 1 | 11 |  |  |
| Gao *et al.* 2017 | Plants | 1 | 5 |  |  |
| Total (N unique) |  | 699 | 2,449,451 | 615 | 2,356,590 |
| Total (N total) |  | 4,028 | 6,383,161 | 3,675 | 6,055,549 |

**Figure 1.**
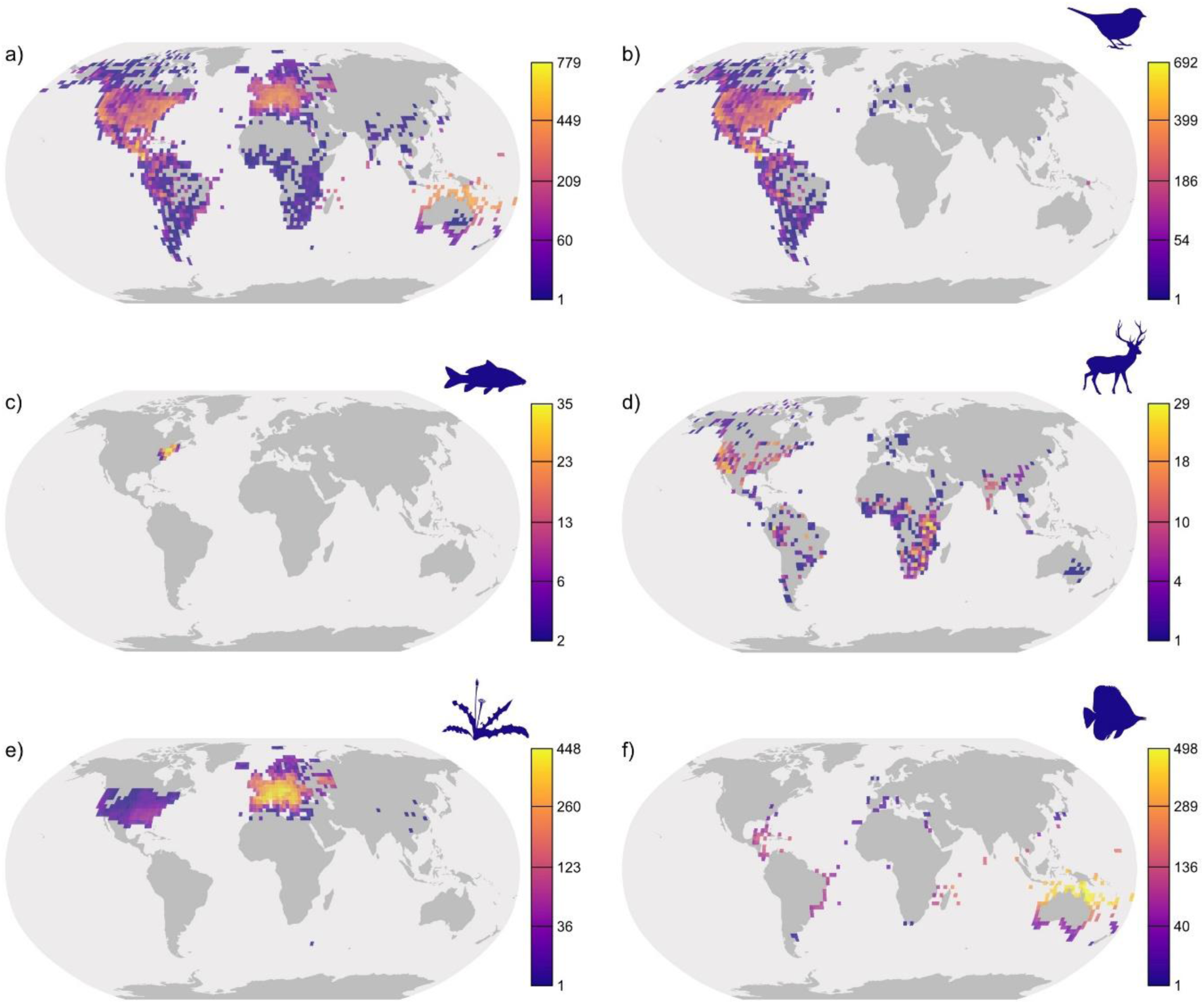
Spatial distribution of 6,383,161 abundance observations for 4,028 species compiled this study: a) all taxonomic groups combined, b) birds, c) freshwater fishes, d) mammals, e) plants and f) reef fishes. Data for 3,060 animal and 615 plant species were subsequently used in the statistical analyses. Maps represent the number of species with abundance data per 2.5° grid cell (for display purposes only), reprojected into the Robinson projection. Values range from dark purple (low number of species) to light yellow (high number species), with breaks obtained using a square-root transformation. Data are compiled from 14 studies conducted across multiple geographic scales.

## RESULTS

### Effects of species traits and range size on abundance–distance relationships

We used meta-analytical mixed-effects models (that accounted account for intra-specific heterogeneity) to summarise the compiled abundance–distance relationships into grand mean and subgroup effect sizes. This enabled us to test for the effects of dispersal-related species traits and range size variables on support for abundant-centre patterns.

Overall, animal species did not conform to abundant-centre patterns (*z* = 1.501, df = 3,059, *P* = 0.133) whereas plant species tended to decrease in abundance further away from their range centroids (*z* = - 16.547, df = 614, *P* < 0.0001), indicating support for abundant-centre patterns (Fig. 2; Fig. 3). Both models showed a significant proportion of unaccounted heterogeneity between abundance–distance slopes (Table 3). Model fail-safe numbers, i.e., the minimum number of non-significant studies that could be included until the grand mean effect becomes non-significant, were high for both animals (N = 668,729) and plants (N = 4,048,392). Inclusion of species traits significantly reduced this proportion of heterogeneity in animal and plant abundance–distance slopes (Table 3). Log likelihood tests indicated significant effects of taxonomic group and range size for animal abundance–distance slopes and significant effects of functional group, life span and range size for plant abundance–distance slopes (Table 3; Fig. 2; Fig. 3).

**Figure 2.**
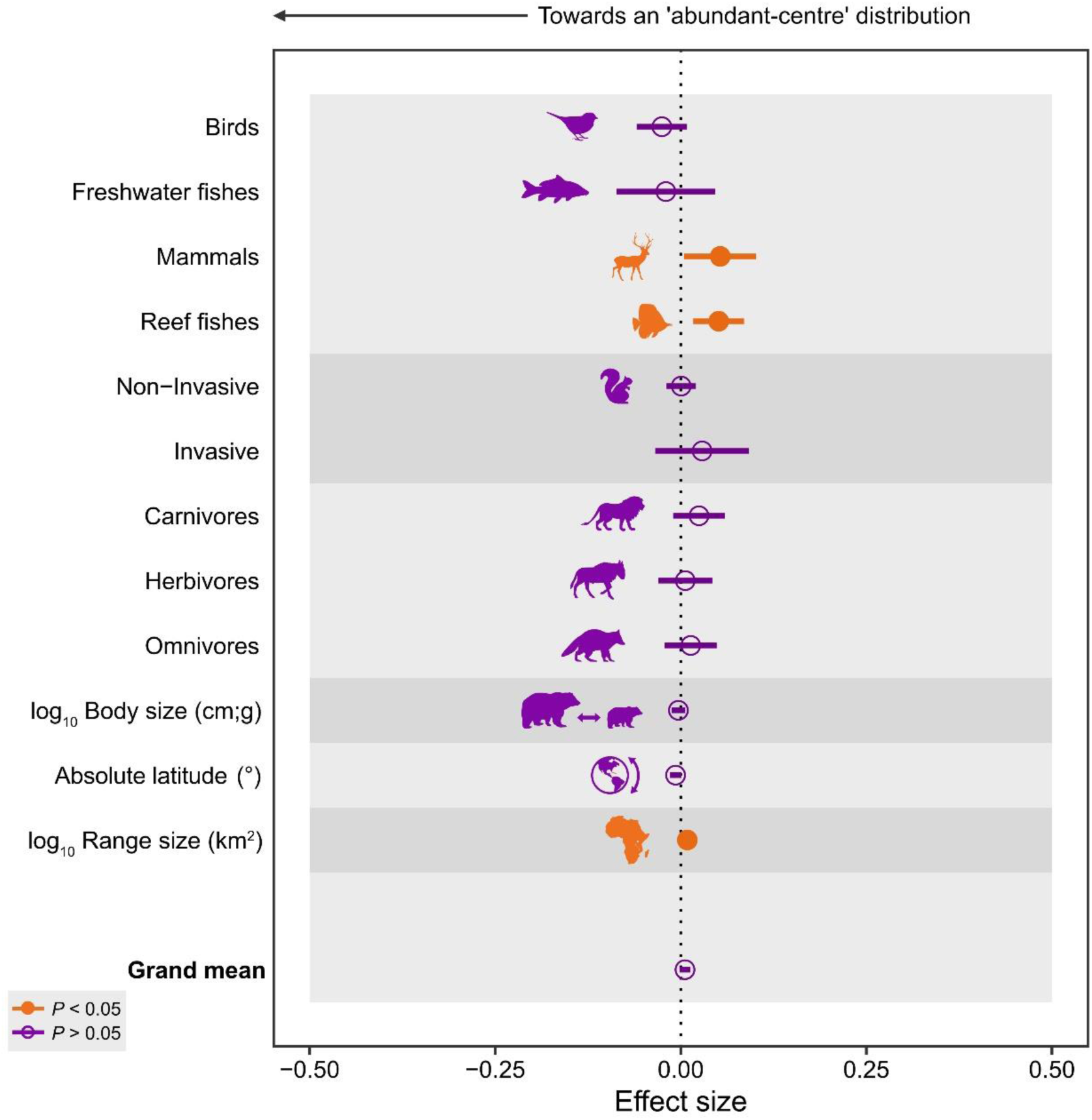
Modelled effects of species traits on abundance–distance slopes (Fisher’s z) for 3,060 animal species, ordered by moderator group. Negative effects (left of dotted line) indicate an ‘abundant-centre’ distribution effect. Effect sizes were derived from marginal mean estimates for categorical moderators (Taxonomic group, Invasiveness and Feeding guild) and from regression slopes for continuous moderators (log_10_ Body size (cm; g), Absolute latitude (°) and log_10_ Range size (km^2^), each scaled to unit variance). Error bars represent approximated 95% confidence intervals.

**Figure 3.**
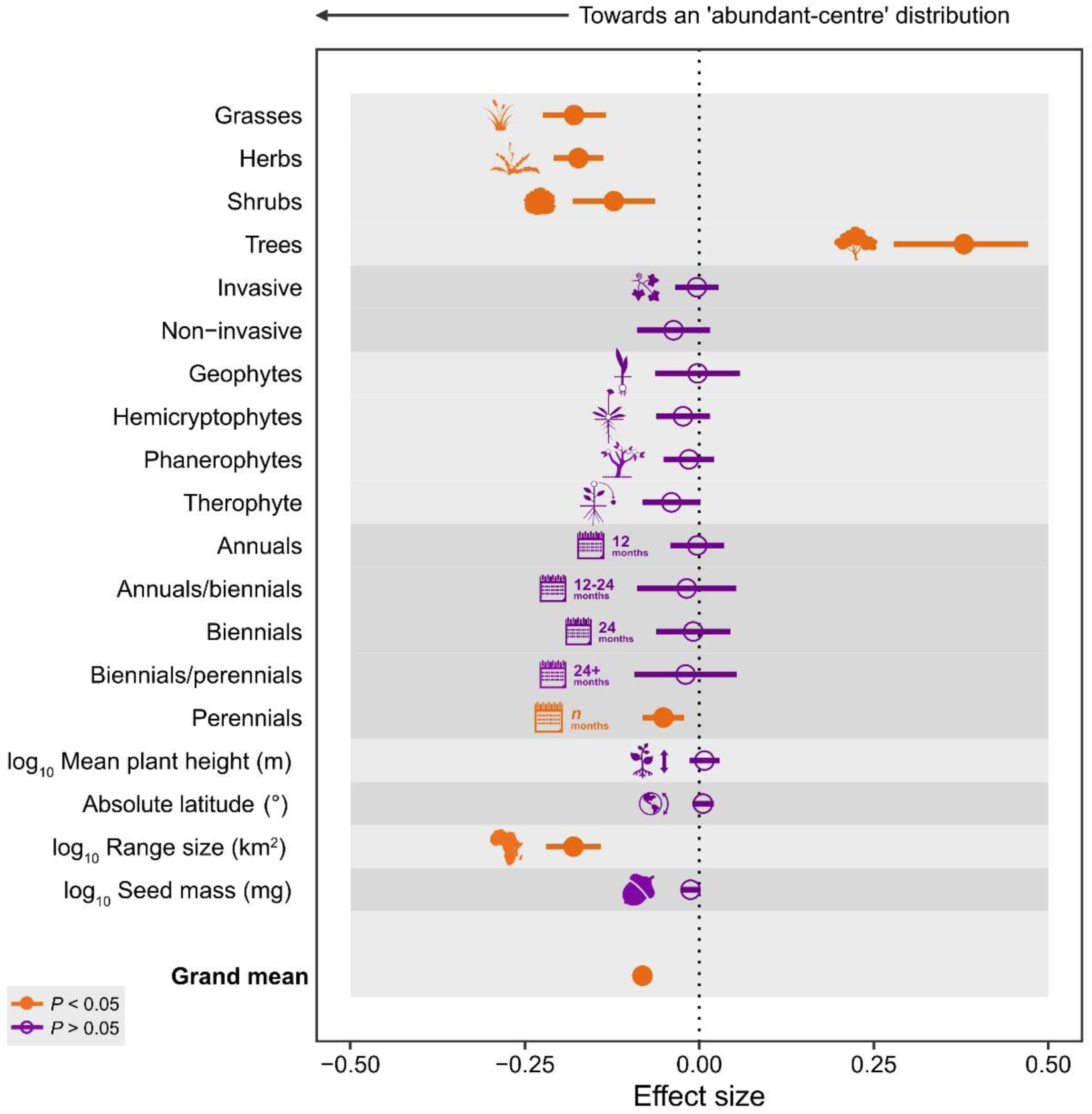
Modelled effects of species traits on abundance–distance slopes (Fisher’s z) for 615 plant species, ordered by moderator group. Negative effects (left of dotted line) indicate an ‘abundant-centre’ distribution effect. Effect sizes were derived from marginal mean estimates for categorical moderators (Functional group, Invasive and Life form) and from regression slopes for continuous moderators (log_10_ Mean plant height (m), Absolute latitude (°), log_10_ Range size (km^2^) and log_10_ Seed mass (mg), each scaled to unit variance). Error bars represent approximated 95% confidence intervals.

For the individual animal species trait analyses, we found positive abundance–distance relationships for mammals and reef fishes (indicating no abundant-centre pattern) (Fig. 2). Range size also showed a positive effect on animal abundance–distance relationships. Invasiveness, feeding guild, body size and latitude were non-significant moderators of animal abundance–distance relationships. For plants, we found that all taxonomic groups showed negative abundance–distance relationships (indicative of abundant-centre patterns), except for trees, which showed positive relationships (Fig. 3). Perennial species and large range sizes were both related to negative abundance–distance relationships while invasiveness, life form, plant height, latitude and seed mass were all non-significant predictors of plant abundance–distance relationships (Fig. 3).

### Interactions between species traits and range size variables

To explore the effects of interactions between combinations of species traits and range size variables on support for abundant-centre patterns, we used a series of meta-analytical mixed-effects interaction models. For animals, there were significant interaction effects between taxonomic group and feeding guild, with herbivorous mammals and reef fishes showing positive abundance–distance relationships (Fig. 4). There were negative interaction effects between taxonomic group and body size, and taxonomic group and latitude. Larger-bodied mammals were more likely to conform to abundant-centre patterns, as were mammals and freshwater fishes from higher latitudes (Fig. 4). For plants, there were significant interaction effects between life form and seed mass, with hemicryptophytes and phanerophytes with larger seeds showing more support for abundant-centre patterns (Fig. 5). Despite the interaction, effect sizes remained positive for hemicryptophytes suggesting no support for abundant-centre patterns in this group. There was also a positive interaction effect between plant height and seed mass suggesting that taller plants with larger seeds showed opposing patterns to the predictions of the abundant-centre hypothesis (Fig. 5).

**Figure 4.**
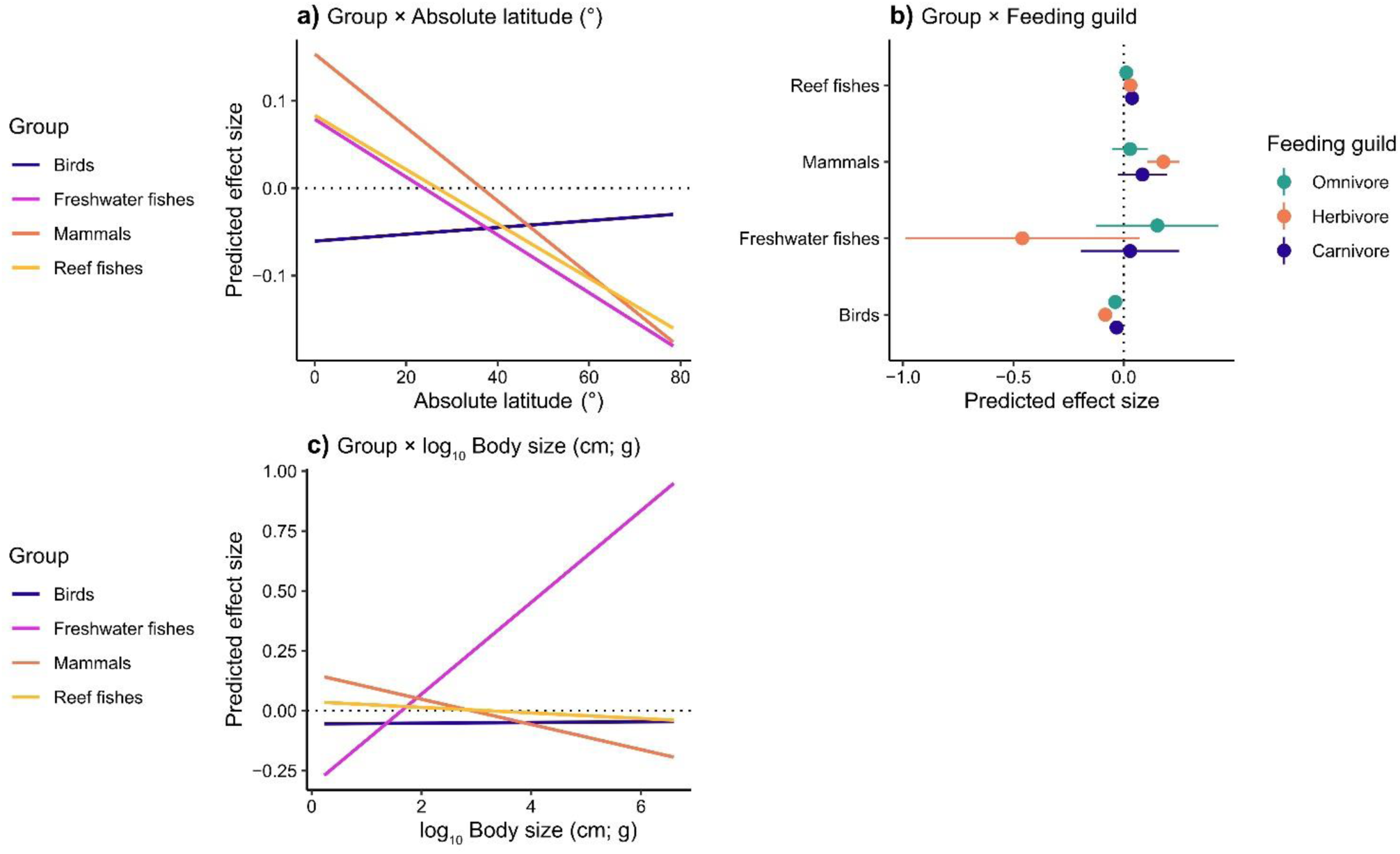
Most important interactions between a) Group × Absolute latitude (°), b) Group × Feeding guild and c) Group × log_10_ Body size (cm; g) used to predict heterogeneity in abundance–distance slopes (Fisher’s z) for 3,060 animal species (Table S4).

**Figure 5.**
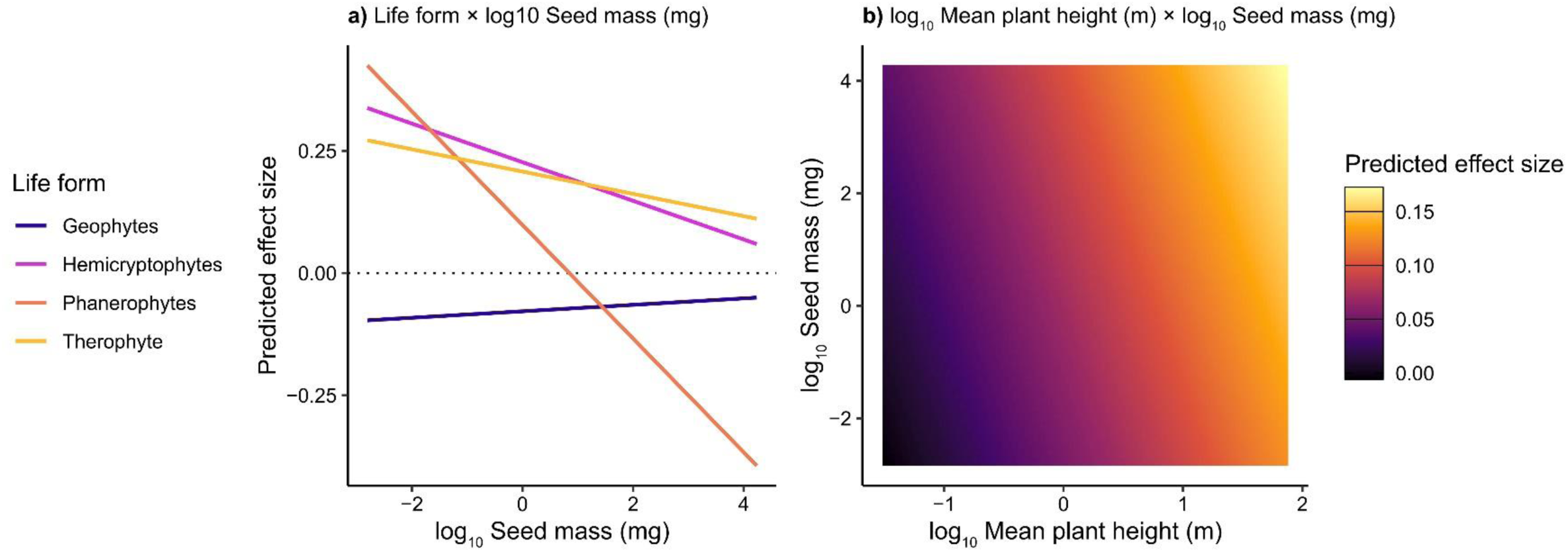
Most important interactions between a) Life form × log_10_ Seed mass (mg) and b) log_10_ Mean plant height (m) × log_10_ Seed mass (mg) used to predict heterogeneity in abundance–distance slopes (Fisher’s z) for 615 plant species (Table S5).

### Quantifying publication bias

There were significant publication year effects for both animals and plants (Fig. S4). More recent animal studies found less support for abundant-centre patterns, whereas more recent plant studies tended to support abundant-centre patterns (Fig. S4). Tests for funnel plot asymmetry were non-significant for both animals (z = - 0.359, P = 0.720) and plants (*z* = -1.329, *P* = 0.184).

## DISCUSSION

Contrary to previous research, which yielded little to no support for the abundant-centre hypothesis (Tuya *et al*. 2008; Hidas *et al*. 2010; Dixon *et al*. 2013; Dallas *et al*. 2017; Santini *et al*. 2018; Ntuli *et al*. 2020), our findings suggest that plants appeared to follow abundant-centre patterns whereas animals tended not to. Moreover, species often conform to abundant-centre patterns when traits related to dispersal capabilities were taken into account. We expected to find conformity to abundant-centre patterns in species with higher dispersal capabilities. This was apparent in plants and larger-bodied mammals, suggesting that species less limited by dispersal constraints are more likely to follow abundant-centre patterns. A recent study found general support for abundant-centre patterns in birds (see Fristoe *et al*. 2022), despite other studies suggesting that abundant-centre patterns are rarely detected in this group (Dallas *et al*. 2017; Santini *et al*. 2019), however we failed to detect these patterns in our bird data. As predicted, invasive animals did not show support for abundant-centre distributions; however, they also did not show significantly positive effects that might be expected in line with invasion theory (Giometto *et al*. 2013).

Unlike other studies, our test of the abundant-centre hypothesis was not restricted geographically. It represents the first global test of the hypothesis compiled from previously published literature. Prior to this study, the largest existing test of the hypothesis found weak support for abundant-centre patterns in geographic and ecological niche space (Dallas *et al*. 2017). They also attempted to explain variation in abundance–distance slopes by exploring the effects of body size, range size and climatic niche area (Dallas *et al*. 2017). We decided not to use the range size variable calculated by Dallas *et al*. (2017), who interpreted minimum convex polygons around sampling points as proxies for a species’ range, and instead sourced the majority of our range size estimates from published IUCN expert range maps (IUCN 2022). Minimum convex polygons have a coarse outer edge at low sampling intensities and are therefore sensitive to outliers (Burgman and Fox 2003), and sampling efforts only cover the majority of a species’ range in exceptional circumstances (such as thorough field surveys of endemic species on small islands; Phiri *et al*. 2015). Furthermore Dallas *et al*. (2017) lends to their use of eBird data to test range-wide abundant-centre patterns in North American birds, which has been regarded as unsuitable when calculating abundance–distance slopes (Fristoe et al. 2022). In an attempt to be as comprehensive as possible, we also included the largest data set on birds from Dallas *et al*. (2017), however, we are aware that caution must be used when interpreting the results because of these known potential flaws. Traits included in Dallas *et al*. (2017), i.e., body size, range size and climatic niche area, explained very little variation within their models (birds: *R^2^* = 3%; trees = *R^2^* = 3%, mammals: *R^2^* = 4% and freshwater fishes: *R^2^* = 2%), whereas our most parsimonious trait interaction models explained 10% of variation for animals and 35% of variation for plants. An explanation for this may simply lie in the number of dispersal-related traits used to test the hypothesis. Dallas *et al*. (2017) also focused on distributions of (Pearson) correlation coefficients when interpreting support for the abundant-centre hypothesis. We explored the distributions of our abundance–distance relationships using Spearman’s rank correlation coefficients which may better account for non-linear relationships between species abundances and distance from range centroids (Fig. S1). This implies that more robust analytical techniques, such as meta-analytical approaches, may be more appropriate when testing broad-scale relationships between species abundances and their environments (see VanDerWal *et al*. 2009; Weber *et al*. 2017; Santini *et al*. 2018).

A study by Ntuli *et al*. (2020) tested the abundant-centre hypothesis on two coastal mussel species, suggesting that abundant-centre patterns are unlikely to be met in species with high dispersal capabilities and instead may be more prominent in species with lower connectivity. Our results suggest the opposite, particularly for plants. One explanation for this may be due to differences in the life histories of the taxa studied as Ntuli *et al*. (2020) concentrated on the genetic implications of the hypothesis in taxa with planktonic life stages (see also Hidas *et al*. 2010; Tam & Scrosati 2011). Unfortunately, in our study we could not compare coastal invertebrates and terrestrial animals because of lack of data.

Accounting for species’ dispersal capabilities when testing the abundant-centre hypothesis enabled strong signals to be detected within a comparatively large data set. Compared to animals, our trait-based method appeared to work better when explaining support for abundant-centre patterns in plants, suggesting that other processes that remain untested here may contribute to the unexplained variation in observed abundance–distance patterns. These biotic and abiotic processes may include, but are not limited to, interspecific interactions (Goldberg and Barton 1992; Robertson 1996), spatiotemporal patterns in resource availability (Theodose and Bowman 1997; Liu *et al*. 2021), climate and environmental suitability (Maitra *et al*. 2022), geodiversity (Bailey *et al*. 2018), and pressures from human activities (Lepczyk *et al*. 2008; Jesse *et al*. 2018). Our focus on species’ dispersal capabilities responded to recent research (Santini *et al*. 2018; Feng & Qiao 2022) which found dispersal to be one of the most important processes driving abundance–distance patterns at a global scale. Therefore, dispersal must be accounted for when testing macroecological hypotheses such as the abundant-centre hypothesis and can be tested using an appropriate combination of dispersal-related species traits.

Similar to other tests of the abundant-centre hypothesis, there are caveats with regard to sampling within our data set. For example, most species’ geographical ranges are likely to have been under-sampled, and abundance estimates are mostly not derived from the full extents of their ranges. Sampling efforts are likely to be biased by differences in data quantity and quality between taxa. For example, bird abundance data are more readily available from open-source repositories (such as eBird – ebird.org) than abundance data for most other taxa. We attempted to account for this by weighting the mixed-effects meta-regression models by species sample size, such that better-sampled species were assigned larger weights in our analyses than less-sampled species. Secondly, there were notable taxonomic gaps within our data set, and a bias towards more commonly sampled groups. During our literature searches, we were unable to obtain any abundance estimates for reptiles, amphibians, marine mammals, pelagic fishes, corals, fungi and most invertebrate groups. Despite these limitations, we were able to calculate abundance–distance slopes for 4,036 species, with 3,675 used in our trait analyses. Thirdly, due to the volume of species data included in this study we were unable to control for spatiotemporal differences between migratory and non-migratory species abundances, which have been shown to influence support for the abundant-centre hypothesis (Osorio-Olvera *et al*. 2020; but see Dallas *et al*. 2017; Chevalier *et al*. 2021). We used IUCN expert range maps, which include a species’ entire geographic range, including both migratory and non-migratory ranges, in an attempt to address this potential limitation. Fourthly, our focus on testing the hypothesis by measure of centrality, as opposed to marginality, may bias our findings. Depending on the shape of a species’ geographic range, such measures can differ largely as occurrences far from the range centroid can also be far from the range edge (Santini *et al*. 2018). Finally, we searched for studies published in English, French and Spanish, but did not obtain any relevant search results in languages other than English which may introduce a bias within our results (see Konno *et al*. 2020). This is likely due to the search terms being optimised for literature published in English.

This study indicates that accounting for dispersal-related traits improves our ability to model abundant-centre patterns across geographic space. Future tests of the abundant-centre hypothesis should aim to (i) conduct more tests on well-known taxonomic groups included here to overcome bias and (ii) include data for under-represented taxonomic groups, e.g., reptiles, amphibians and fungi, to enhance our overall understanding of broad-scale patterns of species’ abundances. Furthermore, the effects of dispersal-related traits should be examined in relation to species’ ecological niche space, in line with recent research (Dallas *et al*. 2017; Osorio-Olvera *et al*. 2019; Dallas and Santini 2020).

Finally, we recommend that future tests of the abundant-centre hypothesis attempt to address other processes, such as interspecific interactions and impacts of human activities, which may underlie abundance–distance relationships and account for additional variation. Our findings contribute to our understanding of how species interact with the wider environment, and have direct implications for invasive species management, population ecology research and will improve our knowledge of how species’ respond to environmental change.

## METHODS

### Literature searches

A systematic literature search was conducted on 23^rd^ July 2021 by querying the ISI Web of Science database (WoS; https://apps.webofknowledge.com/). We used the following search string for an initial broad search: “(abundan* OR abundance-cent* OR abundant niche-cent* OR niche cent* OR abundant-centre hypothesis) AND (range OR geographic range OR range size OR range edge OR species distribution)” using the TITLE field. We retained all studies that 1) comprised peer-reviewed primary studies, 2) presented globally extensive abundance point observations across all taxonomic groups, 3) were published between 1990-2020, 4) included extractable data relating to observed/estimated abundance counts and 5) were published in English, French or Spanish language.

Examination of the returned studies (N = 818) revealed that some key literature was missing from our results. Therefore, we used the studies returned from our initial search as reference sources for additional key search terms to derive an optimised search string (using the *R* package ‘litsearchr’ (Grames *et al*. 2019)). Search terms were extracted from unique study titles, abstracts and tagged keywords (e.g., terms such as ‘range edge’, ‘abundance’ and ‘species range’). We built a keyword co-occurrence network and quantitatively assessed potential search terms using a 60% cumulative cut-off point (see Grames *et al*. 2019). Resulting search terms (N = 326) were grouped into either 1) the ‘species group’, 2) ‘geographic group’ or 3) ‘both groups’ depending on whether the term referred to a species concept or geographic concept. Grouping refers to the string of search terms either side of the Boolean operator ‘AND’. We manually removed irrelevant search terms (N = 268; e.g., ‘field sites’, ‘habitat patch’ and ‘statistically significant’) and retained the most relevant search terms (N = 58) which formed our optimised search string (Table S1). To verify whether our resulting search string was fully optimised, we cross-referenced four key articles that we expected to be included in the optimised search results (Virgós *et al*. 2011; Dixon *et al*. 2013; Baldanzi *et al*. 2013 and Dallas *et al*. 2017), all of which were included. We queried the WoS database using our optimised search string and obtained 531 studies. After screening of titles and abstracts, we retained 23 studies for data extraction.

We supplemented our WoS literature search with the studies included within the foundational synthesis by Sagarin and Gaines (2002; see Table 2). We then conducted a snowball search of the literature that cited Sagarin and Gaines (2002) up until 31^st^ December 2020. This resulted in a literature database of 1,109 studies of which we removed a set of 1,000 studies after screening titles and abstracts and an additional set of 95 studies after screening of full texts, leaving us with an additional 14 studies that were suitable for data extraction.

### Data extraction and processing

From each study and for each species, we extracted raw abundance values and distance from the species’ geographic range centroids (in km). Corresponding authors were contacted via email correspondence if data were not publicly available. Where data were not publicly available and the corresponding author was unable to provide the data, we extracted abundance and distance data from appropriate figures within the published articles using the web-based tool WebPlotDigitizer version 4.5 (Rohatgi 2021; https://automeris.io/WebPlotDigitizer). If abundance data were available but distance values were not, global range maps were obtained in shapefile formats for each species and range centroids were calculated. We obtained global range shapefiles for terrestrial mammals from the IUCN Red List (IUCN 2021; https://www.iucnredlist.org) and for birds from the BirdLife Data Zone version 2020.1 (BirdLife International and the Handbook of the Birds of the World 2020; http://datazone.birdlife.org/home), thus accounting for the entire ranges for migratory and non-migratory species. IUCN range maps have been criticized due to oversimplification of species’ ranges derived from sampling bias (Herkt *et al*. 2017), but represent the most comprehensive spatial data set available for our study species. If global polygon range maps for particularly under-sampled taxonomic groups were unavailable, e.g., invertebrates and some plant species (N spp. = 8), we downloaded species occurrence point data from the Global Biodiversity Information Facility (GBIF; https://www.gbif.org). Occurrence data were then cleaned using the ‘CoordinateCleaner’ R package (Zizka *et al*. 2019) and manual checks were performed to remove any remaining outliers (Zizka *et al*. 2020; Panter *et al*. 2020). We calculated minimum convex hulls for terrestrial species and minimum concave hulls for intertidal species, e.g., *Mytilus* spp. in an attempt to not overestimate their global range sizes by accounting for unsuitable terrestrial environments, which we interpreted as proxies for global species ranges and calculated range centroids using QGIS 3.14.16 (QGIS.org 2022). Then, we calculated the geodesic distances on a sphere (km) between the sampling sites with associated abundance values to obtain the distance to the species’ range centroid, using the WGS84 co-ordinate reference system. Abundance and distance values were log_10_-transformed prior to analyses. Species with unresolved species-level taxonomies (106) as well as species with fewer than five observations were removed from the analysis. We calculated Spearman Rank Correlation Coefficients (*r_s_*) between log_10_(abundance) and log_10_(distance) values (Fig. S1). Negative *r_s_* values are consistent with an ‘abundant-centre’ distribution (Fig. S1).

### Scale effects on abundant–distance relationships

We used three measurements to attempt to explore the effect of scale on ‘abundant-centre’ patterns: 1) we calculated the study extent (km), i.e., the spatial extent at which the study was conducted at encompassing the total study area between sampling locations in the data. Initially we used four categorical levels: ‘Local’ ≤ 250 km, ‘Landscape’ > 250-500 km, ‘Regional’ > 500-1500 km and ‘Continental’ > 1,500 km. 2) We calculated the grain (km^2^), i.e., the spatial scale at which data were collected, which is important because the area of the base unit defines the spatial scale of the study (Field *et al*. 2009), and variation in grain may be reflected in population abundance estimates. Grain was extracted from each study by taking the base unit area for each sampling technique, e.g., sampling units measured in km^2^ (Kallimanis & Koutsias 2012). 3) We calculated study focus (km^2^), defined as the spatial scale at which data were analysed. Often, abundance estimates from individual sampling sites are averaged across larger sampling areas, e.g. a protected area sampled using a number of line transects and multiple sampling points along each transect, with abundance values averaged across all of the sampling points to produce an estimate for each transect. In most cases grain and focus remained the same for each study, e.g. when abundance data were recorded in the form of points within a species’ geographic range. Grain and focus values were log_10_ transformed due to the large variation in the range of these values. We then plotted the distribution of the log_10_- transformed values on separate histograms and visualised the natural breaks in the data. Using these we binned both grain and focus into two new categorical levels: ‘small’ and ‘large’ (-10 to -3 and -3 to 3 on log_10_ scale, respectively). We decided to drop extent from our analyses due to uneven sample sizes: data for only three groups (birds, mammals and plants) global 2,717 species vs. local 146 species. We also dropped focus from our analyses because the natural break categorical bins were identical to those for grain. Grain was subsequently omitted from the statistical analyses due to its strong correlation with animal species group and plant functional group variables, and thus could not be included within the same models.

### Dispersal-related species traits and range size variables

We compiled six traits for animal and eight traits for plant species to examine their effects on abundance–distance relationships (Table 1). Traits were selected based on the morphological, ecological and geographical characteristics of the study species and included: for animals 1) taxonomic group (categorical), 2) body size (continuous), 3) invasiveness (binary 1,0), 4) feeding guild (categorical), 5) range size (km^2^) and 6) absolute latitude (°); and for plants: 1) functional group (categorical), 2) mean plant height (m), 3) seed mass (mg), 4) invasiveness (binary 1,0), 5) life span (categorical), 6) life form (categorical), 7) range size (km^2^) and 8) absolute latitude (°) (see Table 1 for an overview). Due to small sample sizes (N = 9 species), invertebrates were dropped prior to statistical analysis. To examine the effects of body size, we compiled body mass (g) data for mammals and birds, and snout-vent lengths (SVL; cm) for freshwater and reef fishes to produce the trait variable ‘body size’. For plants, we used mean plant height (m) as a proxy for body size. Mean plant height was used instead of maximum plant height as these were the only data available for our selected species, and notable effects of plant height on species abundance patterns would be reflected in either measurement. Where plant height and seed mass data were unavailable, we supplemented our data with gap-filled measurements from Bruelheide *et al*. (2018) which were estimated using Bayesian Hierarchical Probabilistic Matrix Factorization (BHPMF; Schrodt *et al*. 2015). Trait data for plant functional groups were sourced from the BiolFlor database (Kühn *et al*. 2004) and the corresponding levels ‘dwarf shrub’ (N = 11 species) and ‘subshrub’ (N = 4 species) were merged into the level ‘shrub’ to produce four distinct categorical levels: ‘grasses’, ‘herbs’, ‘shrubs’ and ‘trees’. Plant life-form data followed the classification of Raunkiær (1934) but due to small sample sizes for ‘chamaephytes’ (woody plants with perennating buds borne close to the soil surface), we merged these with the ‘phanerophytes’ (woody perennial plants with buds at a distance from the surface, such as trees and shrubs). Invasiveness was assessed using a binary approach (1 = invasive and 0 = non-invasive) according to the Invasive Species Specialist Group’s Global Invasive Species Database (ISSG GISD; http://www.iucngisd.org/gisd/). For both animal and plant species, absolute latitude (°) was calculated as the absolute value of the range centroid. The following species traits were log_10_-transformed prior to analysis: body size (cm; g), mean plant height (m), seed mass (mg) and range size (km^2^) to account for right-skew within the data. We tested for, but did not find, an indication for multicollinearity between continuous species trait variables using a correlation threshold value ≥ 0.70 in the R package ‘hmisc’ (Harrell Jr 2022) and visualised this using ‘pheatmap’ (Kolde 2019) (Fig. S2).

### Meta-analyses and meta-regression

All data analyses were performed in R version 4.0.5 (R Core Team 2021). We used the R package ‘metafor’ (Viechtbauer 2010) to explore the effects of species traits and scale on species abundance–distance relationships. We analysed the animal and plant data separately because of their different sets of traits and the different sampling methods (e.g. abundances estimated from line transects for animals vs. vegetation plot-based cover estimates for plants).

### Grand mean effects and mixed-effects meta-regression models

Rank-correlation estimates between species’ abundances and range-centre distances were transformed to Fisher’s z-scores (to achieve approximate normality, hereafter ‘effect sizes’). In all following analyses, the influence of each effect size was weighted by the inverse of its effect size variance (to down-weight the influence of sparsely sampled species). For animal and plant data, we calculated the grand mean effect size of abundance–distance relationships separately with an intercept-only model that included a random term for each effect size (a common meta-analytical practice to account for anticipated overdispersion in aggregated data). The resulting grand mean effect sizes were determined to be significant if the approximated 95% confidence intervals did not include zero. For each model, we calculated the between-effect size variance (*τ^2^*), unaccounted heterogeneity (Q) and percentage of variability (*I²*).

To explore the effect of dispersal-related species traits and range size variables, we expanded the outlined grand mean effects models to mixed-effects models by adding the proposed species traits and range size moderators as fixed effects. Significance of individual moderators was determined with log likelihood ratio tests between the full model that included all moderators and nested models in which the focal moderator was omitted (both fit with maximum likelihood). From the full models with all moderators, we quantified the impact of continuous moderators using the respective coefficient estimates and quantified the subgroup-level average effect sizes of categorical moderators with estimated marginal mean effects (while keeping all non-focal moderators at their mean value, using the ‘emmeans’ package (Lenth 2021)

### Meta-regression interaction models

To explore potential interaction effects between pairs of moderators, we applied a multimodel inference approach. Separately for animal and plant data, we constructed all mixed-effects models with all potential combinations of the following interaction and their respective fixed effects terms. For animal species, we included interactions between taxonomic group and feeding guild, log_10_ range size (km^2^), log_10_ body mass (cm; g), and absolute latitude (°) and an interaction between feeding guild and log_10_ range size (km^2^). For plant species, we included interactions between life form and log_10_ range size (km^2^), absolute latitude (°) and log_10_ seed mass (mg), interactions between life span and log_10_ range size (km^2^), absolute latitude (°) and log_10_ seed mass (mg), interactions between mean plant height (m) and log_10_ range size (km^2^), absolute latitude (°) and log_10_ seed mass (mg) and an interaction between log_10_ seed mass (mg) and log_10_ range size (km^2^). All competing models were fit with maximum likelihood and ranked according to the Akaike Information Criterion (AIC), which favours the models that explain a high amount of variation with a small number of fixed and interaction effects (using the dredge function of the ‘MuMIn’ package; Barton 2020). The model with the lowest AIC scores (hereafter the ‘most parsimonious model’) was then refit with restricted maximum likelihood to yield correct parameter estimates.

We checked for publication bias within our data using a series of Egger’s regression tests for funnel plot asymmetry, and mixed-effects models with ‘publication year’ fitted as a fixed effect moderator, for both animal and plant species. Finally, for both animals and plants we computed model fail-safe numbers, i.e., the minimum number of non-significant studies that could be added to the data before the grand mean effect becomes non-significant, using the Rosenthal Approach (Rosenthal 1979).

## EXTENDED DATA

**Table S1.** Initial literature search string used to collate relevant literature on the ISI’s Web of Science database (23^rd^ July 2021), with an optimized search string using the R package ‘litsearchr’ (Grames et al. 2019). The initial search returned 818 results and the optimized search returned 1,019 results.

| Initial search string | Optimized search string |
| --- | --- |
| (abundan* OR abundance-cent* OR<br>abundant niche-cent* OR niche cent*<br>OR abundant-centre hypothesis) | ((("abund* centr*" OR "centr* hypothesi*" OR "local*<br>abund*" OR "rang* centr*" OR "rang* edges" OR "rang*<br>margin*" OR "speci* abund*")) |
| **AND** | **AND** |
| (range OR geographic range OR range<br>size OR range edge OR species<br>distribution) | ("distribut* rang*" OR "elev* gradient*" OR "geograph*<br>rang*" OR "rang* edge" OR "rang* limit*" OR "rang*<br>size*" OR "speci* distribut*" OR "speci* rang*" OR "abund*<br>centr*" OR "centr* hypothesi*" OR "elev* rang*" OR "geograph*<br>distribut*" OR "local* abund*" OR "rang* centr*" OR "rang*<br>margin*" OR "spatial* distribut*")) |

**Table S2.** Morphological, ecological and geographical species traits (hereafter ‘moderators’) used to explore the effects on 3,060 animal and 615 plant abundance–distance relationships. Traits presented with their subgroups and units, inclusion within each taxonomic group, and trait data sources.

| Traits/moderators | Subgroups/units | Taxonomic group |  | References |
| --- | --- | --- | --- | --- |
|  |  | Animals | Plants |  |
| Morphological |  |  |  |  |
| Group | birds, freshwater fishes, mammals and reef fishes | ✓ |  | N/A |
| Body size | centimetres; grams (cm; g) | ✓ |  | Froese & Pauly 2022; Tobias *et al.* 2022; Cooke *et al.* 2022 |
| Mean plant height | metres (m) |  | ✓ | Kattge *et al.* (2020) |
| Ecological |  |  |  |  |
| Functional group | grasses, herbs, shrubs and trees |  | ✓ | Kühn *et al.* (2004) |
| Invasiveness | binary (1,0) | ✓ | ✓ | Invasive Species Specialist Group (2015) |
| Life span | annuals, annuals/biennials, biennials, biennials/perennials, perennials |  | ✓ | Kühn *et al.* (2004) |
| Life form | geophytes, therophytes, hemicryptophytes, phanerophytes |  | ✓ | Kühn *et al.* (2004) |
| Seed mass | milligrams (mg) |  | ✓ | Kattge *et al.* (2020) |
| Feeding guild | carnivores, omnivore, herbivores | ✓ |  | Froese & Pauly 2022; Cooke *et al.* 2022 |
| Geographical |  |  |  |  |
| Range size | square kilometres (km<sup>2</sup>) | ✓ | ✓ | Froese & Pauly (2022); Tobias *et al.* (2022); Sporbert *et al.* (2020); (L. Santini, unpublished data); Shalom *et al.* 2020; CDH (E. Welk *et al.*, unpublished data) |
| Absolute latitude | degrees (°) | ✓ | ✓ | Enquist *et al.* (2016); Froese & Pauly (2022); Tobias *et al.* (2022); Cooke *et al.* (2022); Sporbert *et al.* 2020; (L. Santini, unpublished data) |

**Table S3.**
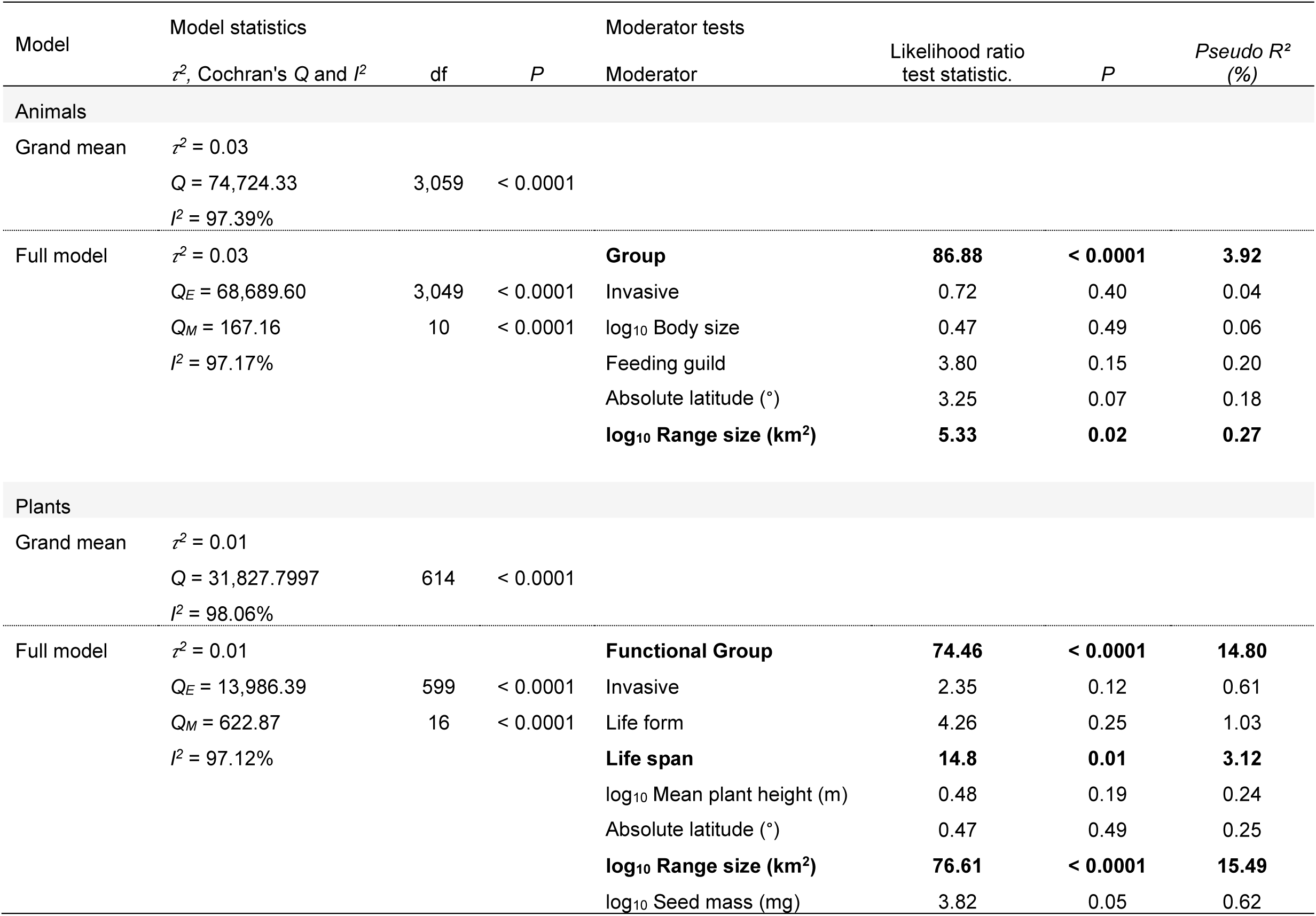
Model results for the grand mean effect and the significance of moderators for abundance– distance slopes (Fishers’ z) for 3,060 animal and 615 plant species. Grand mean models were characterized by the between-species variance (*τ*^2^), unaccounted heterogeneity (Q) and percentage of variability (*I*²). Significance of individual moderators was determined with a log likelihood ratio test between the full model that included all moderators and nested model in which the focal moderator was omitted.

**Table S4.**
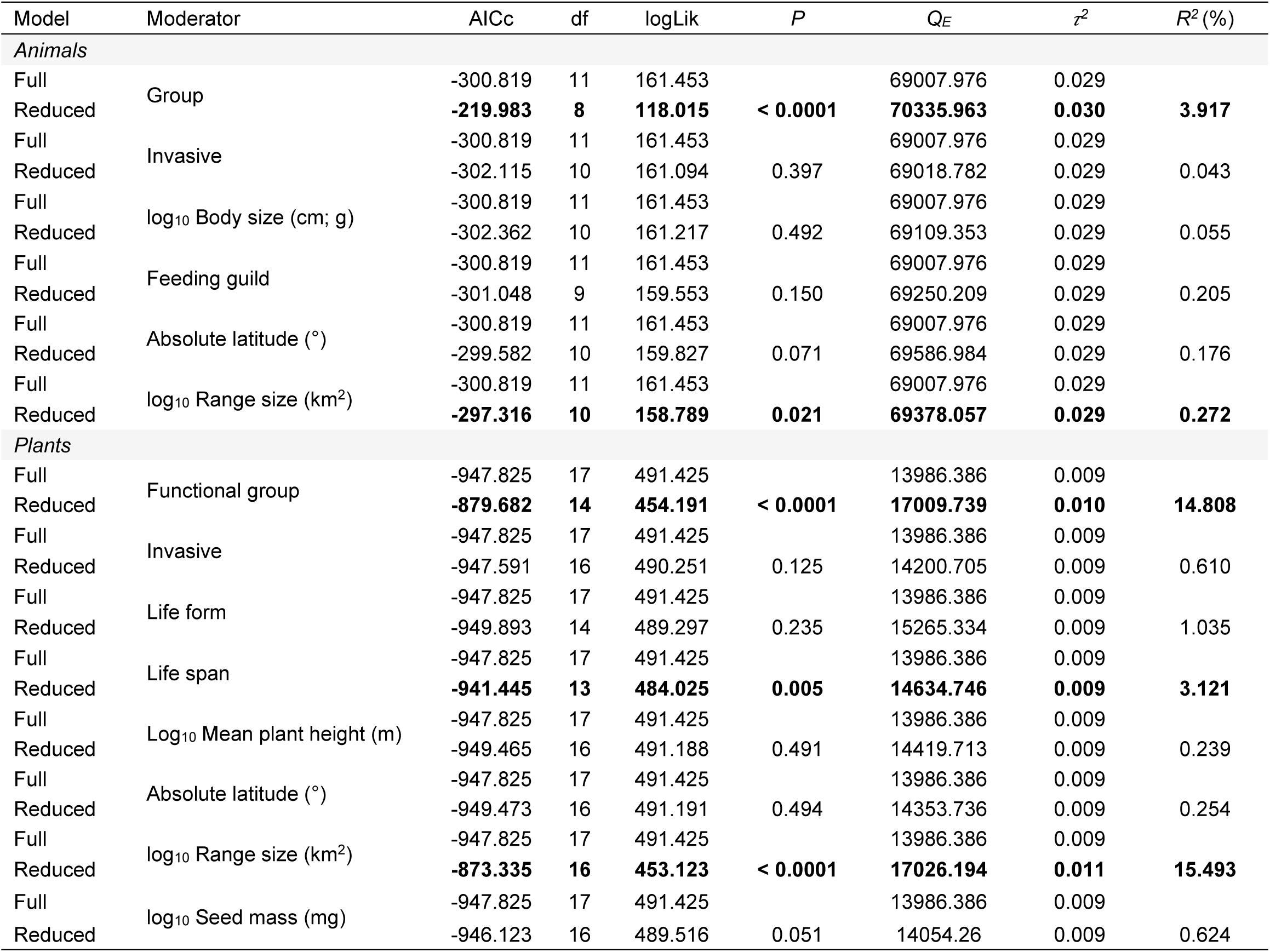
Outputs and proportional variance explained from full and reduced singular moderator meta-regression models, exploring the effects of species traits on abundance–distance relationships for 3,060 animal and 615 plant species. Significant moderators in bold.

**Table S5.**
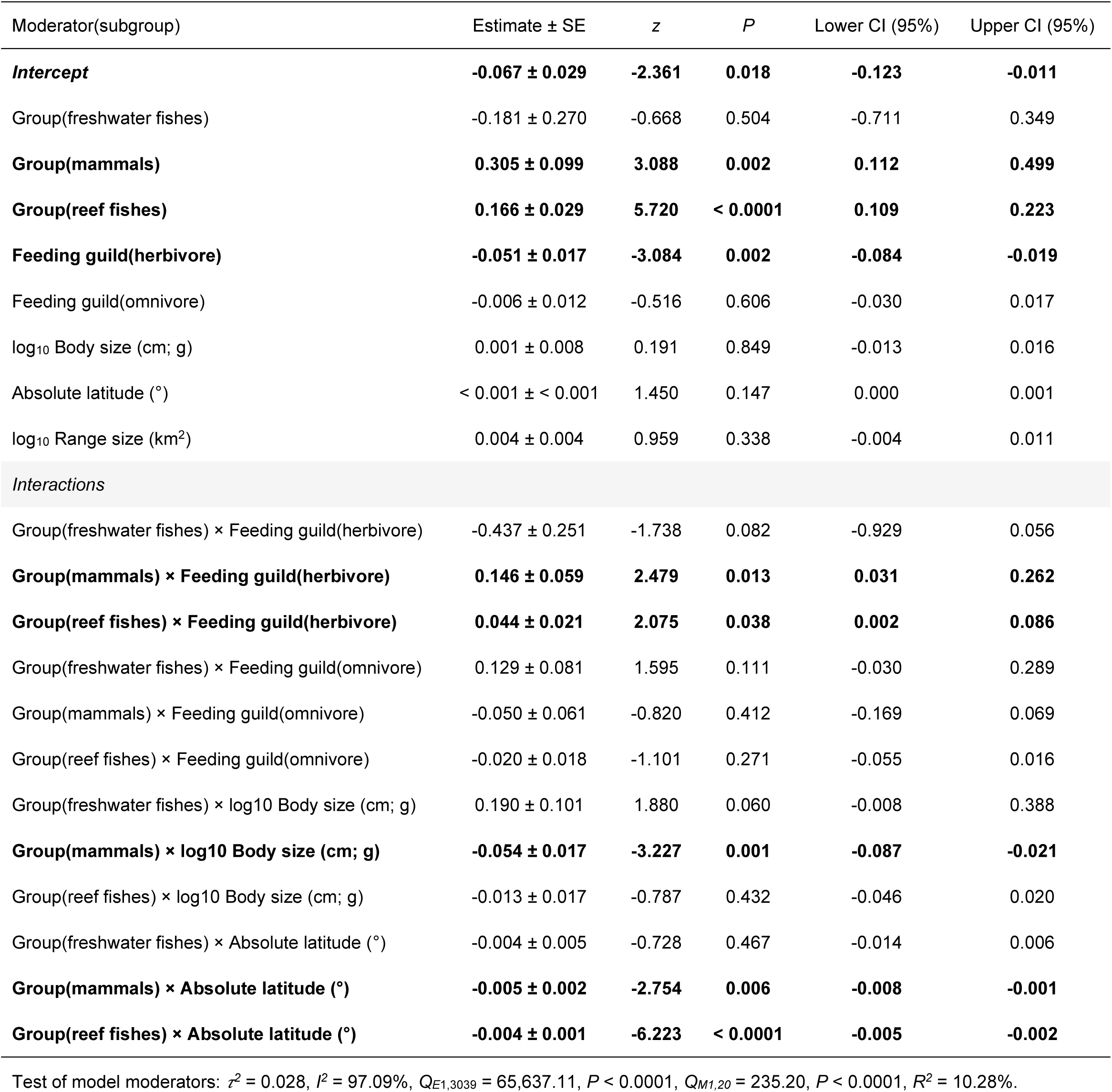
Moderator subgroup effects from the most parsimonious mixed-effects meta-regression interaction model exploring the effects of species traits on 3,060 animal abundance–distance relationships. Significant negative effects are consistent with ‘abundant-centre’ distributions. Significant effects in bold, SE = standard error, CI = confidence interval). Reference subgroups: Group(birds) and Feeding guild(carnivores).

**Table S6.** Moderator subgroup effects from the most parsimonious mixed-effects meta-regression interaction model exploring the effects of species traits on 615 plant abundance–distance relationships. Significant negative effects are consistent with ‘abundant-centre’ distributions. Significant effects in bold, SE = standard error, CI = confidence interval). Reference subgroups: Life form(geophytes) and Life span(annuals).

| Moderator(subgroup) | Estimate ± SE | z | P | Lower CI (95%) | Upper CI (95%) |
| --- | --- | --- | --- | --- | --- |
| *Intercept* | 0.053 ± 0.312 | 0.171 | 0.864 | -0.558 | 0.665 |
| Life form(hemicryptophyte) | 0.305 ± 0.320 | 0.954 | 0.340 | -0.322 | 0.932 |
| Life form(phanerophyte) | 0.177 ± 0.325 | 0.543 | 0.587 | -0.461 | 0.814 |
| Life form(therophyte) | 0.286 ± 0.326 | 0.877 | 0.381 | -0.353 | 0.925 |
| Life span(annual/biennial) | -0.01 ± 0.034 | -0.285 | 0.776 | -0.076 | 0.056 |
| Life span(biennial) | -0.013 ± 0.026 | -0.514 | 0.608 | -0.064 | 0.037 |
| Life span(biennial/perennial) | -0.026 ± 0.036 | -0.737 | 0.461 | -0.096 | 0.044 |
| **Life span(perennial)** | **-0.046 ± 0.014** | **-3.213** | **0.001** | **-0.075** | **-0.018** |
| **log<sub>10</sub> Mean plant height (m)** | **0.039 ± 0.014** | **2.773** | **0.006** | **0.012** | **0.067** |
| log<sub>10</sub> Range size (km<sup>2</sup>) | -0.066 ± 0.042 | -1.555 | 0.120 | -0.149 | 0.017 |
| log<sub>10</sub> Seed mass (mg) | 0.007 ± 0.032 | 0.206 | 0.837 | -0.056 | 0.069 |
| Absolute latitude (°) | 0.006 ± 0.006 | 0.926 | 0.354 | -0.006 | 0.018 |
| *Interactions* |  |  |  |  |  |
| Life form(hemicryptophyte) × log<sub>10</sub> Range size (km<sup>2</sup>) | -0.015 ± 0.044 | -0.343 | 0.732 | -0.102 | 0.071 |
| Life form(phanerophyte) × log<sub>10</sub> Range size (km<sup>2</sup>) | 0.058 ± 0.043 | 1.362 | 0.173 | -0.026 | 0.142 |
| Life form(therophyte) × log<sub>10</sub> Range size (km<sup>2</sup>) | -0.068 ± 0.045 | -1.512 | 0.131 | -0.157 | 0.020 |
| **Life form(hemicryptophyte) × log<sub>10</sub> Seed mass (mg)** | **-0.046 ± 0.019** | **-2.385** | **0.017** | **-0.084** | **-0.008** |
| **Life form(phanerophyte) × log<sub>10</sub> Seed mass (mg)** | **-0.123 ± 0.024** | **-5.166** | **< 0.0001** | **-0.170** | **-0.076** |
| Life form(therophyte) × log<sub>10</sub> Seed mass (mg) | -0.029 ± 0.021 | -1.393 | 0.164 | -0.071 | 0.012 |
| Life form(hemicryptophyte) × Absolute latitude (°) | -0.005 ± 0.006 | -0.820 | 0.413 | -0.018 | 0.007 |
| Life form(phanerophyte) × Absolute latitude (°) | -0.009 ± 0.006 | -1.493 | 0.136 | -0.022 | 0.003 |
| Life form(therophyte) × Absolute latitude (°) | 0.001 ± 0.007 | 0.096 | 0.924 | -0.012 | 0.013 |
| **log<sub>10</sub> Mean plant height (m) × log<sub>10</sub> Seed mass (mg)** | **0.030 ± 0.013** | **2.277** | **0.023** | **0.004** | **0.057** |
| log<sub>10</sub> Range size (km<sup>2</sup>) × log<sub>10</sub> Seed mass (mg) | 0.007 ± 0.004 | 1.643 | 0.100 | -0.001 | 0.015 |

Test of model moderators: $\tau^2 = 0.009$ , $\hat{P} = 96.92\%$ , $Q_{E1,592} = 13,308.83$ , $P < 0.0001$ , $Q_{M1,22} = 289.48$ , $P < 0.0001$ , $R^2 = 35.27\%$ .

**Figure S1.**
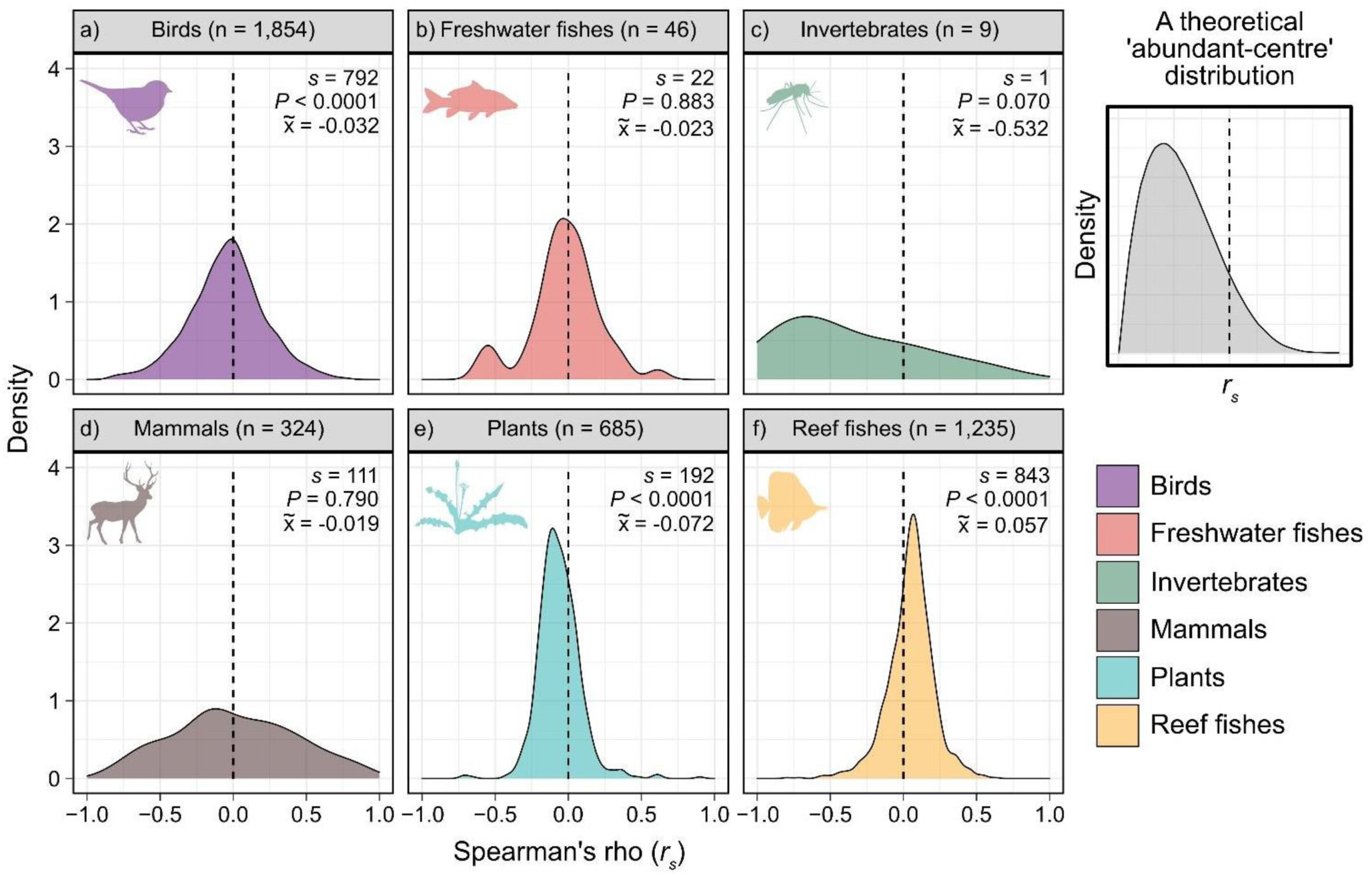
Data exploration phase showing kernel density distributions for Spearman’s Rank Correlation Coefficients (*r_s_*) between log(abundance) and log(distance) by taxonomic group. An ‘abundant-centre’ distribution would be expected to feature predominantly negative *r_s_* values (illustrated in top-right panel). One-sample Sign Tests were used to test for significant differences between median *r_s_* values and zero (*s* = One-sample Sign-Test statistic, x̄ median *r_s_*). Invertebrates were later dropped from the formal analysis due to low sample sizes (N spp. = 9). Species sample sizes are different to those used in the meta-regression models due to data cleaning.

**Figure S2.**
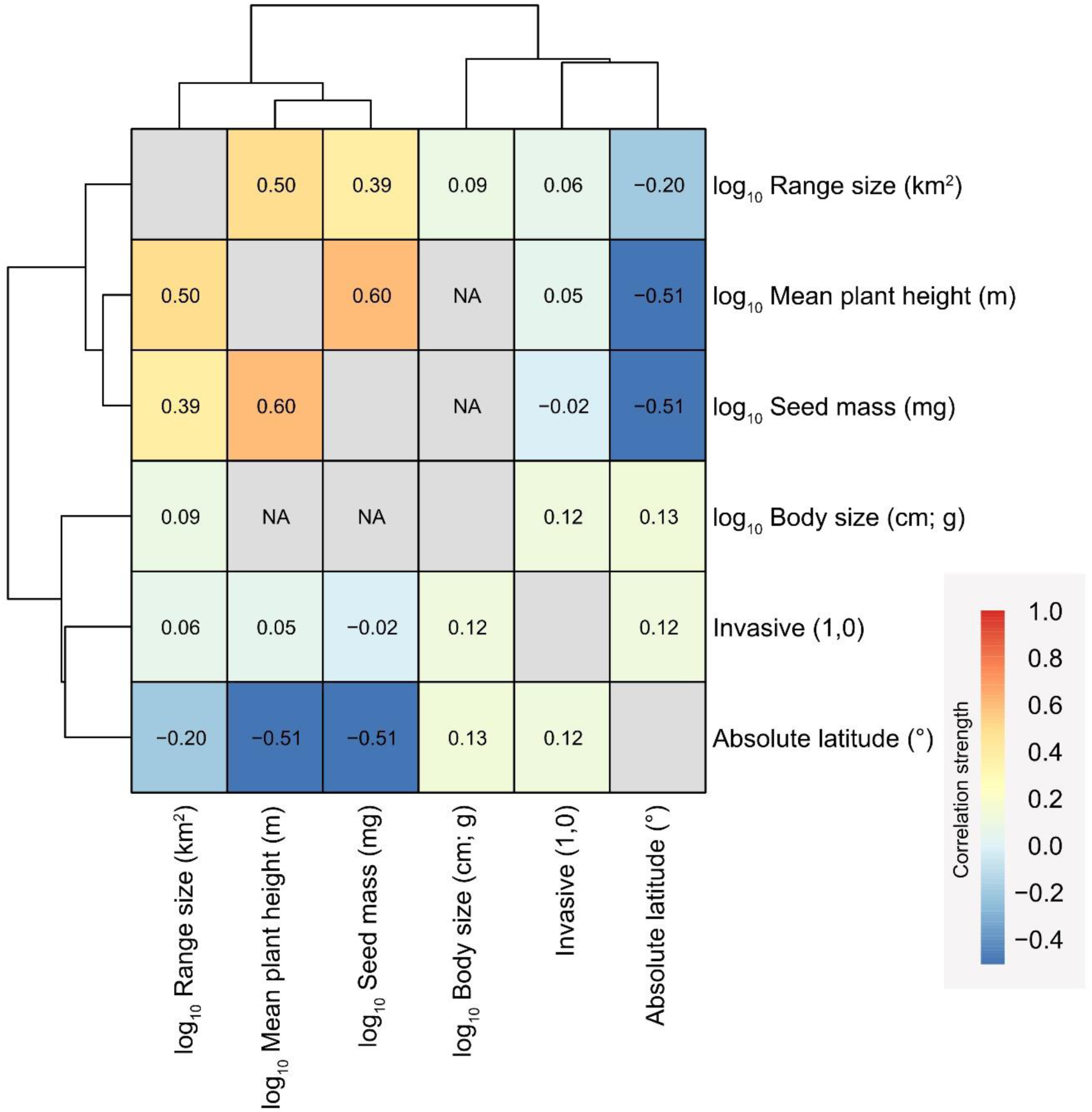
Output from the test of multicollinearity (Spearman Rank Correlation) between continuous species trait variables fitted as moderators in the series of meta-regression models. Continuous moderators were considered ‘colinear’ if the Spearman Rank Correlation coefficient ≥ 0.70.

**Figure S3.**
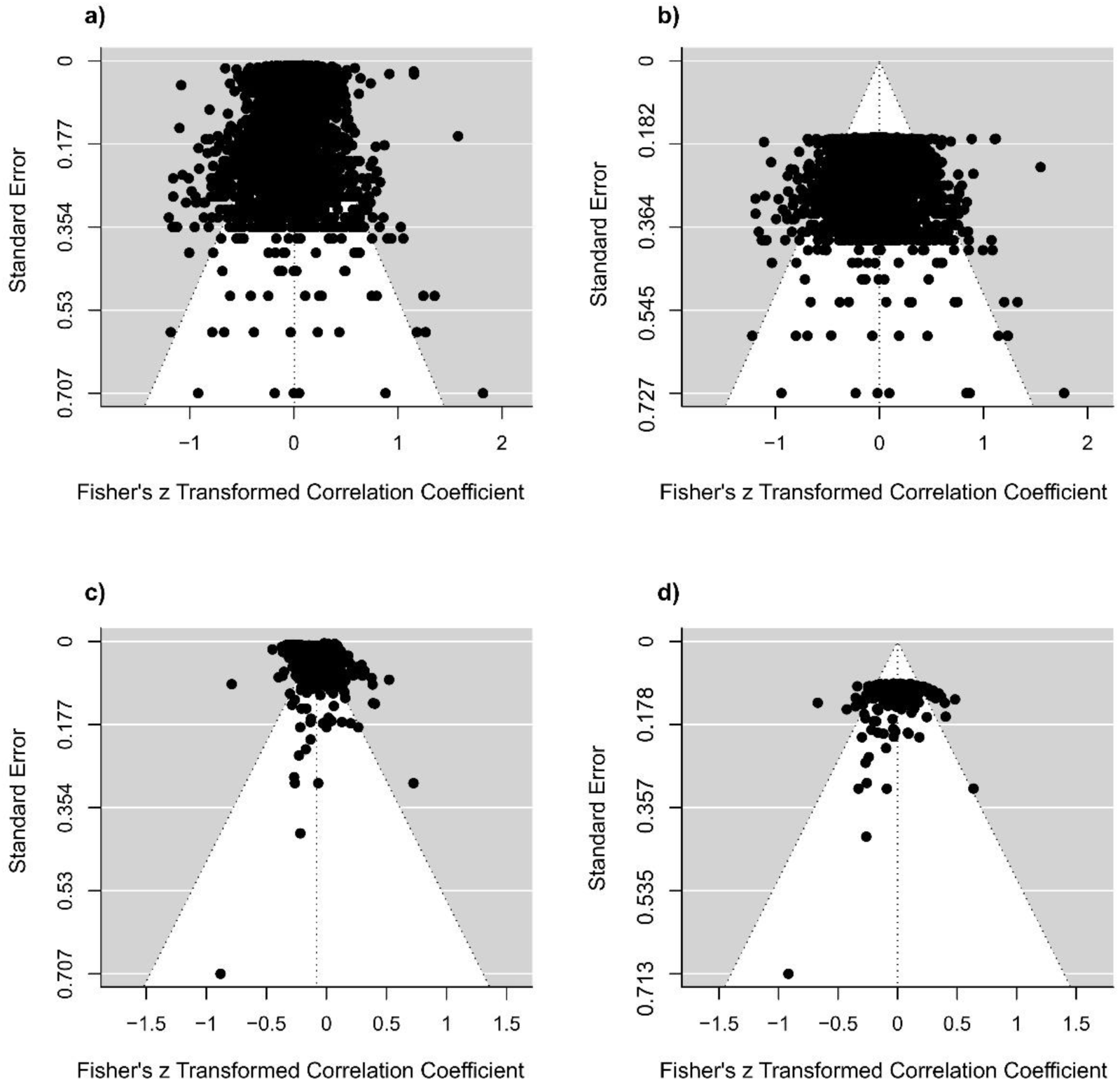
Funnel plots derived from the meta-regression models exploring abundance–distance relationships for 3,060 animal and 615 plant species. Residual outputs from a) animal grand mean model, b) animal full moderator model, c) plant grand mean model and d) plant full moderator model.

**Figure S4.**
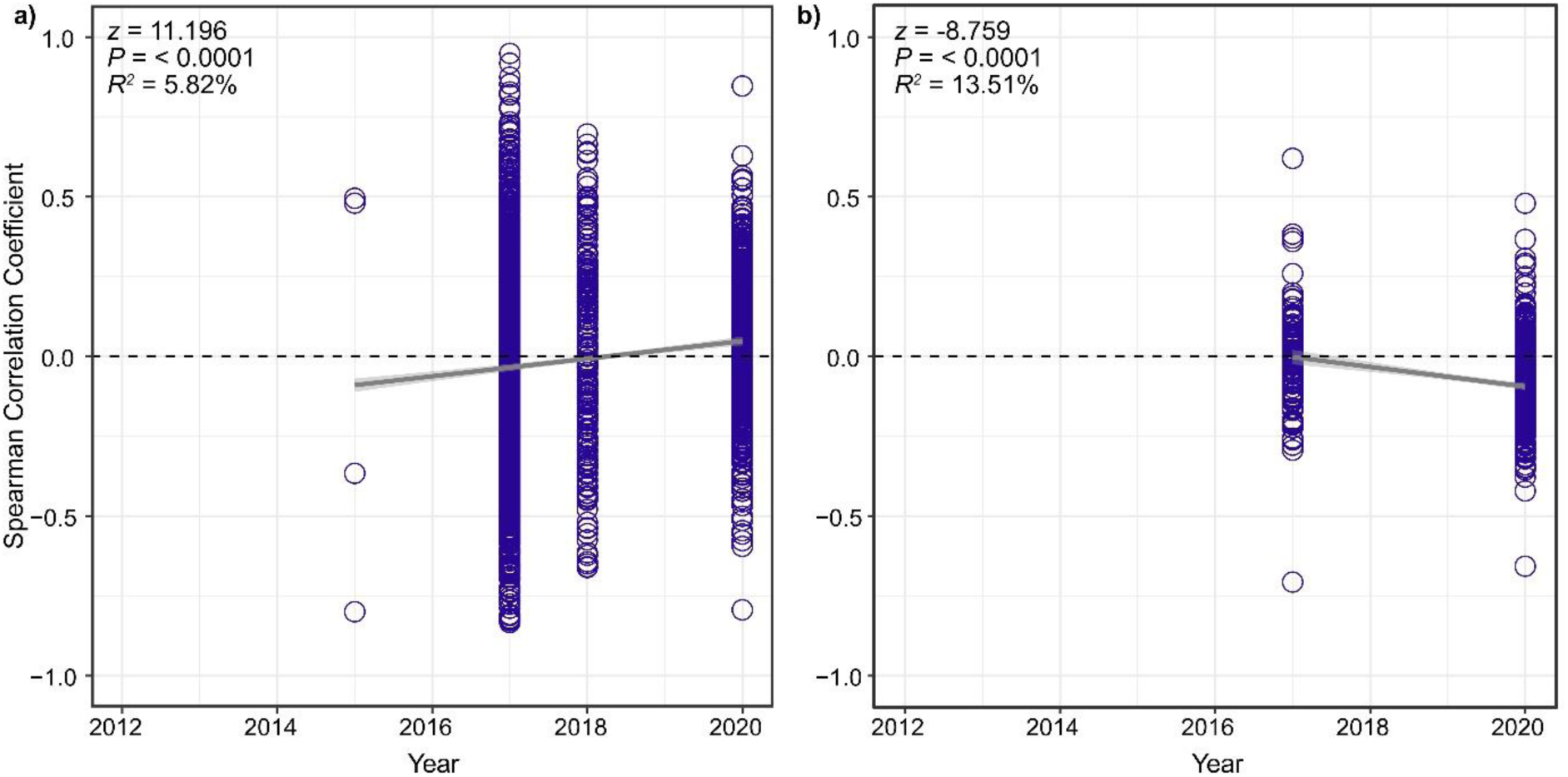
Effects of publication year on abundance–distance slopes for a) 3,060 animal species and b) 615 plant species used in this study. Data sourced from 14 studies (2012–2020). Outputs derived from the animal and plant mixed-effects models, weighted by species sample size, with ‘year’ fitted as the fixed effect moderator. Shading around lines represent 95% confidence intervals.

## REFERENCES

Baer, K.C. & Maron, J.L. 2019. Declining demographic performance and dispersal limitation influence the geographic distribution of the perennials forb *Astragalus utahensis* (Fabaceae). Journal of Ecology 107: 1250–1262.

Bailey, J.J., Boyd, D.S. & Field, R. 2018. Models of upland species’ distributions are improved by accounting for geodiversity. Landscape Ecology 33: 2071–2087.

Baiser, B., Gravel, D., Cirtwill, A.R., Dunne, J.A., Fahimipour, A.K., Gilarranz, L.J., Grochow, J.A., Li, D., Martinez, N.D., McGrew, A., Poisot, T., Romanuk, T.N., Stouffer, D.B., Trotta, L.B., Valdovinos, F.S., Williams, R.J. Wood, S.A. & Yeakel, J.D. 2019. Ecogeographical rules and the macroeocology of food webs. Global Ecology and Biogeography 28: 1204–1218.

Baldanzi, S., McQuaid, C.D., Cannicci, S. & Porri, F. 2013. Environmental domains and range-limiting mechanisms: testing the abundant centre hypothesis using southern African sandhoppers. PLoS ONE 8: e54598.

Barton, K. 2020. MuMIn: multi-model inference. R package version 1.43.17 https://cran.r-project.org/package=MuMIn

Beck, J., Ballesteros-Mejia, L., Buchmann, C.M., Dengler, J., Fritz, S.A., Gruber, B., Hof, C., Jansen, F., Knapp, S., Kreft, H., Schneider, A-K., Winter, M. & Dormann, C.F. 2012. What’s on the horizon for macroecology? Ecography 35: 673–683.

Bergmann, K.G.L.C. 1847. Über die Verhältnisse der wärmeokönomie der Thiere zu ihrer Grösse. Göttinger Studien 3: 595–708.

Blackburn, T.M., Gaston, K.J., Quinn, R.M. & Gregory, R.D. 1999. Do local abundances of British birds change with proximity to range edge? Journal of Biogeography 26: 493–505.

BirdLife International and Handbook of the Birds of the World. 2020. Bird species distribution maps of the world. Version 2020.1. Available at http://datazone.birdlife.org/species/requestdis.

Brown, J.H. 1984. On the relationship between abundance and distribution of species. The American Naturalist 124: 255–279.

Bruelheide, H., Dengler, J., Purschke, O., Lenior, J., Jiménez-Alfaro, B., Hennekens, S.M., Botta-Dukát, Z., Chytrý, M., Field, R., Jansen, F., Kattge, J., Pillar, V.D., Schrodt, F., Mahecha, M.D., Peet, R.K., Sandel, B., van Bodegom, P., Altman, J., Alvarez-Dávila, E., Arfin Khan, M.A.S., Attorre, F., Aubin, I., Baraloto, C., Barroso, J.G., Bauters, M., Bergmeier, E., Biurrun, I., Bjorkman, A.D., Blonder, B., Čarni, A., Cayuela, L., Černý, T., Cornelissen, J.H.C., Craven, D., Dainese, M., Derroire, G., De Sanctis, M., Díaz, S., Doležal, J., Farfan-Rios, W., Feldpausch, T.R., Fenton, N.J., Garnier, E., Guerin, G.R., Gutiérrez, A.G., Haider, S., Hattab, T., Henry, G., Hérault, B., Higuchi, P., Hölzel, N., Homeier, J., Jentsch, A., Jürgens, N., Kącki, Z., Karger, D.N., Kessler, M., Kleyer, M., Knollová, I., Korolyuk, A.Y., Kühn, I., Laughlin, D.C., Lens, F., Loos, J., Louault, F., Lyubenova, M.I., Malhi, Y., Marcenò, C., Mencuccini, M., Müller, J.V., Munzinger, J., Myers-Smith, I.H., Neill, D.A., Miinemets, Ü., Orwin, K.H., Ozinga, W.A., Penuelas, J., Pérez-Haase, A., Petřík, P., Phillips, O.L., Pärtel, M., Reich, P.B., Römermann, C., Rodrigues, A.V., Sabatini, F.M., Sardans, J., Schmidt, M., Seidler, G., Espejo, J.E.S., Silveira, M., Smyth, A., Sporbert, M., Svenning, J-C., Tang, Z., Thomas, R., Tsiripidis, I., Vassilev, K., Violle, C., Virtanen, R., Weiher, E., Welk, E., Wesche, K., Winter, M., Wirth, M. & Jante, U. 2018. Global trait-environment relationships of plant communities. Nature Ecology & Evolution 2: 1906–1917.

Burgman, M.A. & Fox, J.C. 2003. Bias in species range estimates from minimum convex polygons: implications for conservation and options for improved planning. Animal Conservation 6: 19–28.

Burnham, K.P. & Anderson, D.R. 2002. Model selection and multimodel inference: a practical information-theoretic approach, second edition. Springer, New York, New York, USA.

Burner, R.C., Styring, A.R., Rahman, M.A. & Sheldon, F.H. 2019. Occupancy patterns and upper range limits of lowland Bornean birds along an elevation gradient. Journal of Biogeography 46: 2583–2596.

Chaiyes, A., Escobar, L.E., Willcox, E.V., Duengkae, P., Suksavate, W., Watcharaanantapong, P., Pongpattananurak, N., Wacharapluesadee, S. & Hemachudha, T. 2020. An assessment of the niche centroid hypothesis: *Pteropus lylei* (Chiroptera). Ecosphere 11: e03134.

Chevalier, M., Broennimann, O. & Guisan, A. 2021. Using a robust multi-settings inference framework on published datasets still reveals limited support for the abundant centre hypothesis: more testing needed on other datasets. Global Ecology and Biogeography 30: 2211–2228.

Dallas, T. & Santini, L. 2020. The influence of stochasticity, landscape structure and species traits on abundant-centre relationships. Ecography 43: 1341–1351.

Dallas, T., Decker, R.R. & Hastings, A. 2017. Species are not most abundant in the centre of their geographic range or climatic niche. Ecology Letters 20: 1526–1533.

Dallas, T., Santini, L., Decker, R.R. & Hastings, A. 2020. Weighing the evidence for the abundant-center hypothesis. Biodiversity Informatics 15: 81–91.

Dixon, A., Herlihy, C.R. & Busch, J.W. 2013. Demographic and population-genetic test provide mixed support for the abundant centre hypothesis in the endemic plant *Leavenworthia stylosa*. Molecular Ecology 22: 1777–1791.

Enquist B.J., Condit R., Peet R.K., Schildhauer M. & Thiers B.M. 2016. Cyberinfrastructure for an integrated botanical information network to investigate the ecological impacts of global climate change on plant biodiversity. PeerJ Preprints 4: e2615v2.

Ehrlén, J. & Morris, W.F. 2015. Predicting changes in the distribution and abundance of species under environmental change. Ecology Letters 18: 303–314.

Feng, X. & Qiao, H. 2022. Accounting for dispersal using simulated data improves understanding of species abundance patterns. Global Ecology and Biogeography 31: 200–214.

Feldman, R.E., Anderson, M.G., Howerter, D.W. & Murray, D.L. 2015. Where does environmental stochasticity most influence population dynamics? An assessment along a regional core-periphery gradient for prairie breeding ducks. Global Ecology and Biogeography 24: 896–904.

Field, R., Hawkins, B.A., Cornell, H.V., Currie, D.J., Diniz-Filho, A.F., Guégan, F., Kaufman, D.M., Kerr, J.T., Mittelbach, G.G., Oberdorff, T., O’Brien, E.M. & Turner, J.R.G. 2009. Spatial species-richness gradients across scales: a meta-analysis. Journal of Biogeography 36: 132–147.

Freeman, B.G. & Beehler, B.M. 2018. Limited support for the “abundant centre” hypothesis in birds along a tropical elevational gradient: implications for the fate of lowland tropical species in a warmer future. Journal of Biogeography 45: 1884–1895.

Fristoe, T.S., Vilela, B., Brown, J.H. and Botero, C.A. 2022. Abundant-core thinking clarifies exceptions to the abundant-center distribution pattern. Ecography e06365.

Froese, R. and D. Pauly. 2022. FishBase. World Wide Web electronic publication. https://www.fishbase.org version (02/2022).

Gao, W-Q., Ni, Y-Y., Xue, Z-M., Wang, X-F., Kang, F-F., Hu, J., Gao, Z-H., Jiang, Z-P. & Liu, J-F. 2017. Population structure and regeneration dynamics of *Quercus variabilis* along latitudinal and longitudinal gradients. Ecosphere 8: e01737.

Gardner, J.L., Peters, A., Kearney, M.R., Joseph, L. & Heinsohn, R. 2011. Declining body size: a third universal response to warming? Trends in Ecology and Evolution 26: 285–291.

Gaston, K.J., Blackburn, T.M., Greenwood, J.J.D., Gregory, R.D., Quinn, R.M. & Lawton, J.H. 2000. Abundance-occupancy relationships. Journal of Applied Ecology 37: 39–59.

Giometto, A., Rinaldo, A., Carrara, F. & Altermatt, F. 2013. Emerging predictable features of replicated biological invasion fronts. Proceedings of the National Academy of Sciences 111: 297–301.

Goldberg, D.E. & Barton, A.M. 1992. Patterns and consequences of interspecific competition in natural communities: a review of field experiments in plants. The American Naturalist 139: 771–801.

Grames, E.M., Stillman, A.N., Tingley, M.W. & Elphick, C.S. 2019. An automated approach to identifying search terms for systematic reviews using keyword co-occurrence networks. Methods in Ecology and Evolution 10: 1645–1654.

Grinnell, J. 1922. The role of the “accidental”. The Auk 39: 373–380.

Harrell Jr, F. 2022. Hmisc: Harrell Miscellaneous. R package version 4.7.0, https://CRAN.R-project.org/prackage=Hmisc

Herkt, K.M.B., Skidmore, A.K. & Fahr, J. 2017. Macroecological conclusions based on IUCN expert maps: a call for caution. Global Ecology and Biogeography 26: 930–941.

Hidas, E.Z., Ayre, D.J. & Minchinton, T.E. 2010. Patterns of demography for rocky shore, intertidal invertebrates approaching their geographical range limits: tests of the abundant-centre hypothesis in south-eastern Australia. Marine & Freshwater Research 61: 1243–1251.

Hui, C., Richardson, D.M., Robertson, M.P., Wilson, J.R.U. & Yates, C.J. 2011. Macroecology meets invasion ecology: linking the native distributions of Australian acacias to invasiveness. Diversity and Distributions 17: 872–883.

Hutchinson, G.E. 1957. Cold spring harbor symposium on quantitative biology. In Concluding Remarks, pp. 415–427. Cold Spring Harbor Laboratory Press, Long Island, New York.

Invasive Species Specialist Group ISSG. 2015. The Global Invasive Species Database. Version 2015.1 http://www.iucngisd.org/gisd/

IUCN 2022. The IUCN Red List of Threatened Species. Version 2022-1. https://www.iucnredlist.org.

Jesse, W.A.M., Behm, J.E., Helmus, M.R. & Ellers, J. 2018. Human land use promotes the abundance and diversity of exotic species on Caribbean islands. Global Change Biology 24: 4784–4796.

Kallimanis, A.S. & Koutsias, N. 2012. Geographical patterns of corine land cover diversity across Europe: the effect of grain szie and thematic resolution. Progress in Physcial Geography: Earth and Environment 37: 161–177.

Kattge, J., Bönisch, G., Díaz, S. et al. 2020. TRY plant trait database – enhanced coverage and open access. Global Change Biology 26: 119–188.

Kokko, H. & López-Sepulcre, A. 2006. From individual dispersal to species ranges: perspectives for a changing world. Science 313: 789–791.

Kolde, R. 2019. pheatmap: Pretty Heatmaps. R package version 1.0.12, https://CRAN.R-project.org/package=pheatmap

Konno, K., Akasaka, M., Koshida, C., Katayama, N., Osada, N., Spake, R. & Amano, T. 2020. Ignoring non-English-language studies may bias ecological meta-analyses. Ecology and Evolution 10: 6373–6384.

Kühn, I., Durka, W. & Klotz, S. 2004. Biolflor – a new plant-trait database as a tool for plant invasion ecology. Diversity and Distributions 10: 363–365.

Lenth, R.V. 2021. emmeans: Estimated Marginal Means, aka Least-Squares Means. R package version 1.6.2-1. https://CRAN.R-project.org/package=emmeans

Liu, Y., Qi, W., He, D., Xiang, Y., Liu, J., Huang, H., Chen, M. & Tao, J. 2021. Soil resource availability is much more important than soil resource heterogeneity in determining the species diversity and abundance of karst plant communities. Ecology and Evolution 11: 16680–16692.

Lepczyk, C.A., Flather, C.H., Radeloff, V.C., Pidgeon, A.M., Hammer, R.B. & Liu, J. 2008. Human impacts on regional avian diversity and abundance. Conservation Biology 22: 405–416.

Lomolino, M.V., Sad, D.F., Riddle, B.R. & Brown, J.H. 2006. The island rule and research agenda for studying ecogeographical patterns. Journal of Biogeography 33: 1503–1510.

Maitra, A., Pandit, R., Mungee, M. & Athreya, R. 2022. Testing a theoretical framework for the environment-species abundance paradigm: a new approach to the abundant centre hypothesis. bioRxiv https://doi.org/10.1101/2022.01.03.474819

Martínez-Gutiérrez, P.G., Martínez-Meyer, E., Palomares, F. & Fernández, N. 2017. Niche centrality and human influence predict rangewide variation in population abundance of a widespread mammal: the collared peccary (*Pecari tajacu*). Diversity and Distributions 24: 103–115.

Martínez-Meyer, E., Días-Porras, D., Townsend Peterson, A. & Yáñez-Arenas, C. 2013. Ecological niche structure and rangewide abundance patterns of species. Biology Letters 9: 20120637.

Mathys, B.A. & Lockwood, J.L. 2009. Rapid evolution of great kiskadees on Bermuda: an assessment of the island rule to predict the direction of contemporary evolution in exotic vertebrates. Journal of Biogeography 36: 2204–2211.

McGill, B. 2018. The what, how and why of doing macroecology. Global Ecology and Biogeography 28: 6–17.

McGill, B. and C. Collins. 2003. A unified theory for macroecology based on spatial patterns of abundance. Evolutionary Ecology Research 5: 469–492.

McMinn, R.L., Russell, F.L. & Beck, J.B. 2016. Demographic structure and genetic variability throughout the distribution of platte thistle (*Cirsium canescens* Asteraceae). Journal of Biogeography 44: 375–385.

Millien, V., Kathleen Lyons, S., Olson, L., Smith, F.A., Wilson, A.B. & Yom-Tov, Y. 2006. Ecotypic variation in the context of global climate change: revisiting the rules. Ecology Letters 9: 853–869.

Ntuli, N.N., Nicastro, K.R., Zardi, G.I., Assis, J., McQuaid, C.D. & Teske, P.R. 2020. Rejection of the genetic implications of the “abundant centre hypothesis” in marine mussels. Scientific Reports 10: 604.

Osorio-Olvera, L., Yáñez-Arenas, C., Martínez-Meyer, E. & Townsend Peterson, A. 2020. Relationships between population densities and niche-centroid distances in North American birds. Ecology Letters 23: 555–564.

Osorio-Olvera, L., Soberón, J. & Falconi, M. 2019. On population abundance and niche structure. Ecography 42: 1415–1425.

Osorio-Olvera, L., Falconi, M. & Soberón, J. 2016. Sobre la relación entre idoneidad del hábitat y la abundancia poblacional bajo diferentes escenarios de dispersión. Revista Mexicana de Biodiversidad 87: 1080–1088.

Panter, C.T., Clegg, R.L., Moat, J., Bachman, S.P., Klitgård, B.B. & White, R.L. 2020. To clean or not to clean: cleaning open-source data improves extinction risk assessments for threatened plant species. Conservation Science and Practice 2: e311.

Pearce, J. & Ferrier, S. 2001. The practical value of modelling relative abundance of species for regional conservation planning: a case study. Biological Conservation 98: 33–43.

Pérez-Collazos, E., Sanchez-Gómez, P., Jiménez, J.F. & Catalán, P. 2009. The phygeographical history of the Iberian steppe plant *Ferula loscosii* (Apiaceae): a test of the abundant-centre hypothesis. Molecular Ecology 18: 848–861.

Phiri, E.E., McGeoch, M.A. & Chown, S.L. 2015. The abundance structure of *Azorella selago* Hook. f. on sub-Antarctic Marion Island: testing the peak and tail hypothesis. Polar Biology 38: 1881–1890.

Pianka, E.R. 1966. Latitudinal gradients in species diversity: a review of concepts. The American Naturalist 100: 33–46.

Pironon, S., Papuga, G., Villellas, J., Angert, A.L., García, M.B. & Thompson, J.D. 2016. Geographic variation in genetic and demographic performance: new insights from an old biogeographic paradigm. Biological Reviews 92: 1877–1909.

Pironon, S., Villellas, J., Thuiller, W., Eckhart, V.M., Geber, M.A. Moeller, D.A. & García, M.B. 2017. The ‘Hutchinsonian niche’ as an assemblage of demographic niches: implications for species geographic ranges. Ecography 41: 1103–1113.

QGIS.org. 2022. QGIS Geographic Information System. QGIS Association. http://www.qgis.org

R Core Team. 2021. R: A language and environment for statistical computing. R Foundation for Statistical Computing, Vienna, Austria. URL https://www.R-project.org/.

Robertson, D.R. 1996. Interspecific competition controls abundance and habitat use of territorial Caribbean damselfishes. Ecology 77: 885–899.

Raunkiær, C. 1934. The life forms of plants and statistical plant geography, being the collected papers of C. Raunkiær. Clarendon Press.

Rivadeneira, M.M., Hernáez, P., Baeza, J.A., Boltaña, S., Cifuentes, M., Correa, C., Cuevas, A., del Valle, E., Hinojosa, I., Ulrich, N., Valdivia, N., Vásquez, N., Zander, A. and M. Thiel. 2010. Testing the abundant-centre hypothesis using intertidal porcelain crabs along the Chilean coast: linking abundance and life-history variation. Journal of Biogeography 37: 486–498.

Rohatgi, A. 2021. Webplotdigitizer: Version 4.5 https://automeris.io/WebPlotDigitizer

Rosenthal, R. 1979. The file drawer problem and tolerance for null results. Psychological Bulletin 86: 638–641.

Sagarin, R.D. & Gaines, S.D. 2002a. The ‘abundant centre’ distribution: to what extent is it a biogeographical rule? Ecology Letters 5: 137–147.

Sagarin, R.D. & Gaines, S.D. 2002b. Geographical abundance distributions of coastal invertebrates: using one-dimensional ranges to test biogeographic hypotheses. Journal of Biogeography 29: 985–997.

Santini, L., Antão, L.H., Jung, M., Benítez-López, A., Rapacciuolo, G., Di Marco, M., Jones, F.A.M., Haghkerdar, J.M. & González-Suárez, M. 2021. The interface between macroecology and conservation: existing links and untapped opportunities. Frontiers of Biogeography 13.4: e53025.

Santini, L., Pironon, S., Maiorano, L. & Thuiller, W. 2018. Addressing common pitfalls does not provide more support to geographical and ecological abundant-centre hypotheses. Ecography 42: 696–705.

Schrodt, F., Kattge, J., Shan, H., Fazayeli, F., Joswig, J., Banerjee, A. et al. 2015. BHPMF – a hierarchical Bayesian approach to gap-filling and trait prediction for macroecology and functional biogeography. Global Ecology and Biogeography 24: 1510–1521.

Scrosati, R.A. & Freeman, M.J. 2019. Density of intertidal barnacles along their full elevational range of distribution conforms to the abundant-centre hypothesis. PeerJ 7: e6719.

Sexton, J.P., McIntyre, P.J., Angert, A.L. and K.J. Rice. 2009. Evolution and ecology of species range limits. Annual Review of Ecology, Evolution and Systematics 40: 415–436.

Shade, A., Dunn, R.R., Blowes, S.A., Keil, P., Bohannan, B.J.M., Herrmann, M., Küsel, K., Lennon, J.T., Sanders, N.J., Storch, D. & Chase, J. 2018. Macroecology to unite all life, large and small. Trends in Ecology & Evolution 33: 731–744.

Shalom, H.Y., Granot, I., Blowes, S.A., Friedlander, A., Mellin, C., Ferreira, C.E.L., Arias-González, J.E., Kulbicki, M., Floeter, S.R., Chabanet, P., Parravicini, V. & Belmaker, J. 2020. A closer examination of the ‘abundant centre’ hypothesis for reef fishes. Journal of Biogeography 47: 2194–2209.

Soberón, J., Townsend Peterson, A. & Osorio-Olvera, L. 2018. A comment on “species are not most abundant in the centre of their geographic range or climatic niche”. Rethinking Ecology 3: 13–18.

Sporbert, M., Keil, P., Seidler, G., Bruelheide, H., Jandt, U., Aćić, S., Biurrun, I., Campos, J.A., Čarni, A., Chytrý, M., Ćušterevska, R., Dengler, J., Golub, V., Jansen, F., Kuzemko, A., Lenoir, J., Marcenò, C., Moeslund, J.E., Pérez-Haase, A., Rūsiņa, S., Šilc, U., Tsiripidris, I., Vandvik, V., Vasilev, K., Virtanen, R. & Welk, E. 2020. Testing macroecological abundance patterns: the relationship between local abundance and range size, range position and climatic suitability among European vascular plants. Journal of Biogeography 47: 2210–2222.

Stevens, G.C. 1989. The latitudinal gradient in geographical range: how so many species coexist in the tropics. The American Naturalist 133: 240–256.

Stevens, G.C. 1992. The elevational gradient in altitudinal range: an extension of Rapoport’s latitudinal rule to altitude. The American Naturalist 140: 893–911.

Tam, J.C. & Scrosati, R.A. 2011. Mussel and dogwhelk distribution along the north-west Atlantic coast: testing predications derived from the abundant-centre model. Journal of Biogeography 38: 1536–1545.

Theodose, T.A. & Bowman, W.D. 1997. Nutrient availability, plant abundance, and species diversity in two alpine tundra communities. Ecology 78: 1861–1872.

Thompson, P.L. & Gonzalez, A. 2017. Dispersal governs the reorganization of ecological networks under environmental change. Nature Ecology & Evolution 1: 0162.

Tobias, J.A., Sheard, C., Pigot, A.L., Devenish, A.J.M., Yang, J., Sayol, F., Neate-Clegg, M.H.C., Alioravainen, N., Weeks, T.L., Barber, R.A., et al. 2022. AVONET: morphological, ecological and geographical data for all birds. Ecology Letters 25: 581–597.

Trumbo, D.R., Epstein, B., Hohenlohe, P.A. Alford, R.A. Schwarzkopf, L. & Storfer, A. 2016. Mixed population genomics support for the central marginal hypothesis across the invasive range of the cane toad (*Rhinella marina*) in Australia. Molecular Ecology 25: 4161–4176.

Tuya, F., Wernberg, T. & Thomsen, M.S. 2008. Testing the ‘abundant centre’ hypothesis on endemic reef fishes in south-western Australia. Marine Ecology Progress Series 372: 225–230.

VanDerWal, K., Shoo, L.P., Johnson, C.N. & Williams, S.E. 2009. Abundance and the environmental niche: environmental suitability estimated from niche models predicts the upper limit of local abundance. The American Naturalist 174: 283–291.

Viechtbauer, W. 2010. Conducting meta-analyses in R with the metafor package. Journal of Statistical Software 36: 1–48.

Virgós, E., Kowalczyk, R., Trua, A., de Marinis, A., Mangas, J.G., Barea-Azcón, J.M. & Geffen, E. 2011. Body size clines in the European badger and the abundant centre hypothesis. Journal of Biogeography 38: 1546–1556.

Waldock, C., Stuart-Smith, R.D., Edgar, G.J., Bird, T.J. & Bates, A.E. 2019. The shape of abundance distributions across temperature gradients in reef fishes. Ecology Letters 22: 685–696.

Weber, M.M., Stevens, R.D., Diniz-Filho, J.A.F. & Grelle, C.E.V. 2017. Is there a correlation between abundance and environmental suitability derived from ecological niche modelling? A meta-analysis. Ecography 40: 817–828.

Wen, Z., Ge, D., Feijó, A., Du, Y., Cheng, J., Sun, J., Wang, Y. & Xia, L. 2020. Varying support for abundance-centre and congeneric-competition hypotheses along elevation transects of mammals. Journal of Biogeography 48: 616–627.

Yáñez-Arenas, C., Martín, G., Osorio-Olvera, L., Escobar-Luján, J., Castaño-Quintero, S., Chiappa-Carrara, X. & Martínez-Meyer, E. 2020. The abundant niche-centroid hypothesis: key points about unfilled niches and the potential use of supraspecific modeling units. Biodiversity Informatics 15: 92–102.

Zizka, A., Carvalho, F.A., Calvente, A., Baez-Lizarazo, M.R., Cabral, A., Coelho, J.F.R., Colli-Silva, M., Fantinati, M.R., Fernandes, M.F., Ferreira-Araujo, T., Moreira, F.G.L., da Cunha Santos, N.M., Santos, T.A.B., dos Santos-Costa, R.C., Serrano, F.C., da Silva, A.P.A., de Souza, G.C., Tomaz, E.C., de Souza Soares, A., Vale, V.F., Vieira, T.L. & Antonelli, A. 2020. No one-size-fits-all solution to clean GBIF. PeerJ 8: e9916.

Zizka, A., Silvestro, D., Andermann, T., Azevedo, J., Ritter, C.D., Edler, D., Farooq, H., Herdean, A., Ariza, M., Scharn, R., Svantesson, S., Wengström, N., Zizka, V. & Antonelli, A. 2019. CoordinateCleaner: standardized cleaning of occurrence records from biological collection databases. Methods in Ecology and Evolution 10: 744–751.

